# Plant BCL-Domain Homologues play a conserved role in SWI/SNF complex stability

**DOI:** 10.1101/2024.09.17.612632

**Authors:** Joan Candela-Ferre, Jaime Pérez-Alemany, Borja Diego-Martin, Vijaya Pandey, James A. Wohlschlegel, Jorge Lozano-Juste, Javier Gallego-Bartolomé

**Affiliations:** Instituto de Biología Molecular y Celular de Plantas (IBMCP), CSIC-Universitat Politècnica de València, Valencia, 46022, Spain; Department of Biological Chemistry, David Geffen School of Medicine, University of California, Los Angeles, CA, 90095, USA

**Keywords:** SWI/SNF, chromatin, remodeling, Arabidopsis

## Abstract

The SWItch/Sucrose Non-Fermenting (SWI/SNF) complexes are evolutionarily conserved, ATP-dependent chromatin remodelers crucial for multiple nuclear functions in eukaryotes. Recently, plant BCL-Domain Homolog (BDH) proteins were identified as shared subunits of all plant SWI/SNF complexes, significantly impacting chromatin accessibility and various developmental processes in Arabidopsis. In this study, we performed a comprehensive characterization of *bdh* mutants, revealing a previously overlooked impact on hypocotyl cell elongation. Through detailed analysis of BDH domains, we identified a plant-specific N-terminal domain that facilitates the interaction between BDH and the rest of the complex. Additionally, we uncovered the critical role of the BDH β-hairpin domain, which is phylogenetically related to metazoan BCL7 SWI/SNF subunits. While phylogenetic analyses did not identify BDH/BCL7 orthologs in fungi, structure prediction modeling demonstrated strong similarities between the SWI/SNF catalytic modules of plants, animals, and fungi, and revealed the yeast Rtt102 protein as a structural homolog of BDH and BCL7. This finding is supported by the ability of Rtt102 to interact with the Arabidopsis catalytic module subunit ARP7 and partially rescue the *bdh* mutant phenotypes. Further experiments revealed that BDH promotes the stability of the ARP4-ARP7 heterodimer, leading to the partial destabilization of ARP4 in the SWI/SNF complexes. In summary, our study unveils the molecular function of BDH proteins in plant SWI/SNF complexes and suggests that β-hairpin-containing proteins are evolutionarily conserved subunits crucial for ARP heterodimer stability and SWI/SNF activity across eukaryotes.

## Introduction

Chromatin, the association of DNA with histones, enables strong genome compaction in eukaryotic nuclei and participates in numerous nuclear processes such as transcription, replication, repair, and recombination. The basic unit of chromatin is the nucleosome, where 147 base pairs of DNA are wrapped around a histone octamer composed of H2A-H2B dimers and an H3-H4 tetramer (1, 2). These nucleosomes facilitate chromatin compaction and reduce accessibility for DNA-binding proteins, thereby influencing their ability to reach their targets(3). Additionally, nucleosomes can incorporate various histone post-translational modifications and histone variants, fine-tuning their properties and acting as key nuclear signaling elements recognized by diverse histone readers, which transduce this information into specific outputs (4). Therefore, controlling nucleosome positions and modifications is central to most nuclear processes, making their remodeling essential for precise genome function.

Among the various regulators of chromatin remodeling, ATP-dependent chromatin remodelers are critical for controlling nucleosome positioning and occupancy (5). These complexes utilize the energy from ATP hydrolysis to alter the interaction between DNA and histones, participating in nucleosome maturation, sliding, ejection, and composition (5). Consequently, their activity directly influences DNA accessibility, impacting diverse nuclear processes that require a DNA template. The SWItch/Sucrose Non-Fermenting (SWI/SNF) remodelers are an evolutionarily conserved family that play key roles in multiple processes, including the regulation of responses to internal and external signals and cell differentiation (6, 7). In plants, SWI/SNF complexes participate in multiple nuclear processes, such as transcription, DNA repair, gene silencing, and responses to diverse signaling pathways (6, 8). Consequently, mutants in SWI/SNF components exhibit strong developmental defects, impaired growth, and altered stress responses (8–10).

The SWI/SNF family forms modular complexes with multiple subunits, assembled into functionally distinct configurations known as subclasses, defined by signature subunits (6, 11, 12). In yeast, two main subclasses have been identified: RSC and SWI/SNF, while in mammals, there are three distinct assemblies known as BAF, PBAF, and ncBAF (5, 6, 13). Recent studies in plants have revealed the composition of three distinct subclasses: BRM-associated SWI/SNF complex (BAS), SYD-associated SWI/SNF complex (SAS), and MINU-associated SWI/SNF complex (MAS) (9, 10, 12, 14). All these subclasses evolved from a common ancestral SWI/SNF form (12) to meet the specific requirements of various organisms, highlighting their adaptive functions across a diverse range of species. Despite this subclass divergence, a common feature of all SWI/SNF complexes is their catalytic module, which incorporates a Snf2-like SWI/SNF ATPase and a heterodimer of ACTIN (ACT) and/or ACTIN-RELATED PROTEINs (ARP), which interact with the helicase-SANT-associated (HSA) domain of the ATPase (Snf2^HSA^) (11, 15–17). Although the ATPase alone can remodel chromatin in vitro, interaction with an ACT-ARP heterodimer significantly enhances its remodeling activity (18). Notably, this ACT-ARP-Snf2^HSA^ assembly is conserved in other ATP-dependent chromatin remodelers like INO80 and SWR1 and the histone acetyltransferase NuA4 (19, 20). Additionally, in animals, the catalytic module of SWI/SNF complexes includes an additional subunit known as B-cell lymphoma/leukemia protein 7 (BCL7), which is strongly implicated in cancer and is thought to assist the activity of the complex, possibly through its interaction with the ATPase and the nucleosome (11, 21–23). On the other hand, fungi incorporate the Regulator of Ty transposition protein 102 (Rtt102) protein into their SWI/SNF-RSC catalytic modules, which interacts with the the Arp7/Arp9 heterodimer and promotes its compaction, enhancing the remodeling activity of the complex (17, 18, 24). However, no phylogenetic relationship has been found between animal BCL7 and fungal Rtt102, which are considered kingdom specific. Moreover, to our knowledge no functional connection between them has been identified. More recently, immunoprecipitation followed by mass spectrometry (IP-MS) experiments identified the BCL-Domain Homolog (BDH) proteins (aka BCL7) in plants (9, 10, 12, 25). This was shown to interact with the catalytic module of the BAS complex, together with BRM, ARP4, and ARP7, likely through interaction with the BRM ATPase (26, 27). Furthermore, *bdh* mutants were shown to alter plant development and chromatin accessibility over hundreds of genes (9, 26, 27). BDH proteins are conserved across plants and contain an evolutionary conserved domain found in animal BCL7 proteins, suggesting functional conservation (12). No orthologs of BCL7 or BDH have been found in fungi (12), suggesting their loss upon divergence of animals and fungi. Apart from this conserved region, no other domains have been reported in BDH.

In this study, we performed a comprehensive characterization of BDH proteins in Arabidopsis, uncovering an evolutionary connection with animal and fungal SWI/SNF subunits. We thoroughly analyzed *bdh* mutants, revealing an overlooked role of BDH proteins as negative regulators of cell elongation in hypocotyls. Furthermore, we conducted a detailed functional analysis of conserved BDH domains, identifying two important domains for BDH interaction with the complex. Structural modeling revealed a strong connection between plant BDH, animal BCL7, and fungal Rtt102, despite the lack of sequence conservation of the latest. The functional conservation of Rtt102 and BDH was further supported by the Rtt102 ability to interact with Arabidopsis ARP7 and partially complement *bdh* mutants. In vivo studies showed that BDH stabilizes the ARP4-ARP7 heterodimer, resulting in a partial loss of ARP4 protein in the BDH-depleted SWI/SNF complexes, providing a mechanistic explanation for the phenotypic and molecular defects observed in the absence of BDH.

## Results

### BDH affects plant development and hypocotyl elongation

The Arabidopsis genome contains two *BDH* paralogs, *BDH1* and *BDH2*, and T-DNA insertion mutants have been recently identified for these genes (9, 26). Consistent with recent reports, *bdh1* and *bdh2* single mutants had a wild-type (WT) appearance whereas double *bdh1/bdh2* mutants (from now on *bdh* mutants) exhibited significant developmental defects, indicating functional redundancy between BDH proteins in Arabidopsis (Supplementary Figure 1A). The *bdh* mutants presented smaller rosettes with curled leaves and an early flowering phenotype in both long- and short-day conditions (Supplementary Figure 1A-E, Supplementary Data 1). Additionally, the *bdh* mutants formed flowers with an increased number of petals compared to WT plants, although this occurred with low frequency (Supplemental Figure 1F). Furthermore, these mutants had shorter siliques that produced fewer seeds, with the severity of this phenotype varying, as siliques of different sizes were found on the same stem (Supplementary Figure 1G,H, Supplementary Data 1).

We also identified a previously overlooked phenotype in *bdh* mutants, which presented longer hypocotyls in different light conditions (continuous light, long-, and short-days) (Figure 1A, Supplementary Data 2), as well as under a gradient of light intensities and qualities (white light and blue, far-red, and red monochromatic lights) (Supplementary Figure 2A-D, Supplementary Data 3). Furthermore, etiolated hypocotyls, which germinate and grow in darkness, also showed more elongated hypocotyls, suggesting that BDH affects hypocotyl elongation mechanisms independent of light (Figure 1A, Supplementary Figure 2A-D, Supplementary Data 2,3). Measurements of hypocotyl cell numbers revealed no difference between WT and *bdh* mutants, indicating that the longer hypocotyl phenotype resulted from enhanced hypocotyl cell elongation rather than cell division (Supplementary Figure 2E, Supplementary Data 3). To further gain information about the impact of BDH on etiolated hypocotyl growth, we performed a time-course experiment analyzing the growth of WT and *bdh* etiolated hypocotyls during the first 10 days after germination in the dark. Results showed that the *bdh* hypocotyls grew at a same speed than WT until the 4th day after which they continued growing at a faster rate for 2 additional days (Figure 1B, Supplementary Data 2). To shed light into the processes altered in the *bdh* mutant that led to increased hypocotyl elongation, we performed an RNA-seq experiments comparing 5-day-old WT and *bdh* etiolated seedlings. Differential expression analyses indicated hundreds of differentially expressed genes (DEGs) that were up- or down-regulated (1100 and 856, respectively, log2 Fold Change 0.58, q value 0.05) (Figure 1C, Supplementary Data 4). An analysis of enriched gene ontologies (GO) revealed several categories overrepresented in *bdh* mutants like “Suberin biosynthesis” and “Glucosinolate/glycosinolate/S-glycoside catabolism” (Supplementary Figure 3A, Supplementary Data 5). Interestingly, the “Xyloglucan metabolic process” category was also highly enriched due to the up-regulation of 14 *Xyloglucan endotransglucosylase/hydrolases genes* (*XTHs*) in the *bdh* mutant (Figure 1D, Supplementary Figure 3B, Supplementary Data 5,6). XTHs are important enzymes controlling the arrangement of the cell wall and play an important role promoting cell elongation (28) and, therefore, their general up-regulation could explain the *bdh* elongated hypocotyls in the dark. Since this phenotype was also found in light-grown *bdh* seedlings, we searched for up-regulation of *XTH* genes in a published RNA-seq comparing light-grown *bdh* and WT seedlings (9) (Supplementary Data 7). Notably, 9 *XTH* genes were also up-regulated in this condition being 5 commonly up-regulated in dark and light (Figure 1D, Supplementary Data 6). A visual inspection of a published BDH1 and BDH2 ChIP-seq in light-grown seedlings (10) indicated that these shared *XTHs* are direct SWI/SNF targets (Supplementary Figure 3C). Interestingly, the overall overlap of DEGs in the light and dark RNA-seq experiments was not very high indicating that BDHs are generally mobilizing distinct transcriptomes in the different growth conditions (Supplementary Figure 3D, Supplementary Data 8,9). In summary, these results suggest that BDHs act as negative regulators of hypocotyl elongation possibly through the repression of *XTH* genes. Future studies will further investigate the functional connection between SWI/SNF complexes and *XTH* expression.

**Figure 1.**
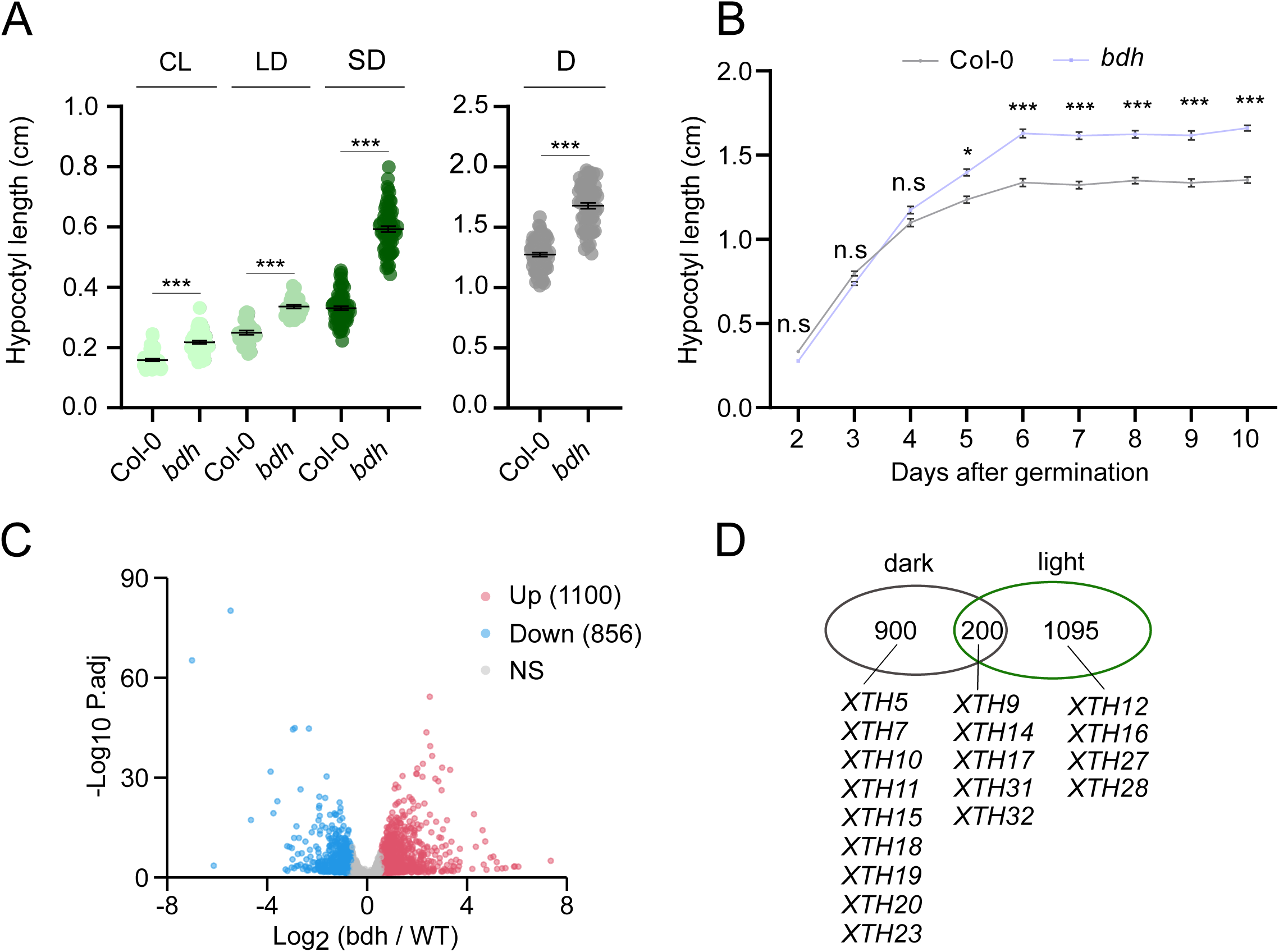
BDH affects plant development and hypocotyl elongation. (A) Hypocotyl length measurements of 7-day-old Col-0 and bdh under Continuous Light (CL), 16h light/8h dark (LD), 8h light/16h dark (SD) and darkness (D) conditions (n=28-63). Error bars represent Mean ± SEM. Asterisks denote statistically significant differences between mean values, as assessed by Student’s t-test (*P < 0.05, **P < 0.01, ***P < 0.001). Two independent biological replicates were conducted with similar results. (B) Time course analysis of hypocotyl length in Col-0 and *bdh* mutant in darkness. Error bars represent Mean ± SEM, n= 42-64. Two independent biological replicates were conducted with similar results. Student’s t-test (*P < 0.05, ***P < 0.001, n.s., not significant). (C) Volcano plot showing differential expression between 5-day-old *bdh* and WT seedlings grown in darkness determined by RNA-seq. Genes showing an adjusted p-value lesser than 0.05 and an absolute fold change over log2 0.58 were considered as DEGs. D. Venn Diagram showing the overlap between *bdh* DEGs detected in darkness (this work) and light^9^ depicting the XTH genes present in each set and the overlap between them.

### Functional dissection of conserved BDH domains

BDH proteins were initially identified through a series of IP-MS experiments in plants using distinct SWI/SNF subunits as baits (9, 12, 14, 25, 26). However, a comprehensive analysis of BDH domain composition and their function remains unreported. Recently, we have reported the evolutionary conservation of BDH based on the sequence homology of a BDH domain, here named BCL domain, with a region in the mammalian SWI/SNF subunit BCL7 (12) (Figure 2A, Supplementary Figure 4A). Structural prediction analyses using AlphaFold (29) for Arabidopsis BDH and Human BCL7 indicated the formation of two antiparallel β-sheets (β-hairpin) in these conserved regions (Figure 2B, Supplementary Figure 4B). Furthermore, detailed phylogenetic analysis across multiple plant species revealed a conserved domain in the most amino-terminal (N-term) region of BDH proteins, which we named N domain (Supplementary Figure 5, Supplementary Data 10). However, structural modeling did not reveal any specific fold in this region (Figure 2B, Supplementary Figure 4B). Interestingly, despite the lack of sequence conservation, structural modeling predicted a conserved downstream alpha-helix fold across multiple plant species, named Alpha domain (Figure 2B, Supplementary Figure 4B,C). Downstream of this domain, sequence conservation is very low even among paralogs (Supplementary Figure 5). An analysis using DISOPRED3 (30) predicted that the identified domains are ordered regions within overall disordered domains in BDH1 (Supplementary Figure 4D).

**Figure 2.**
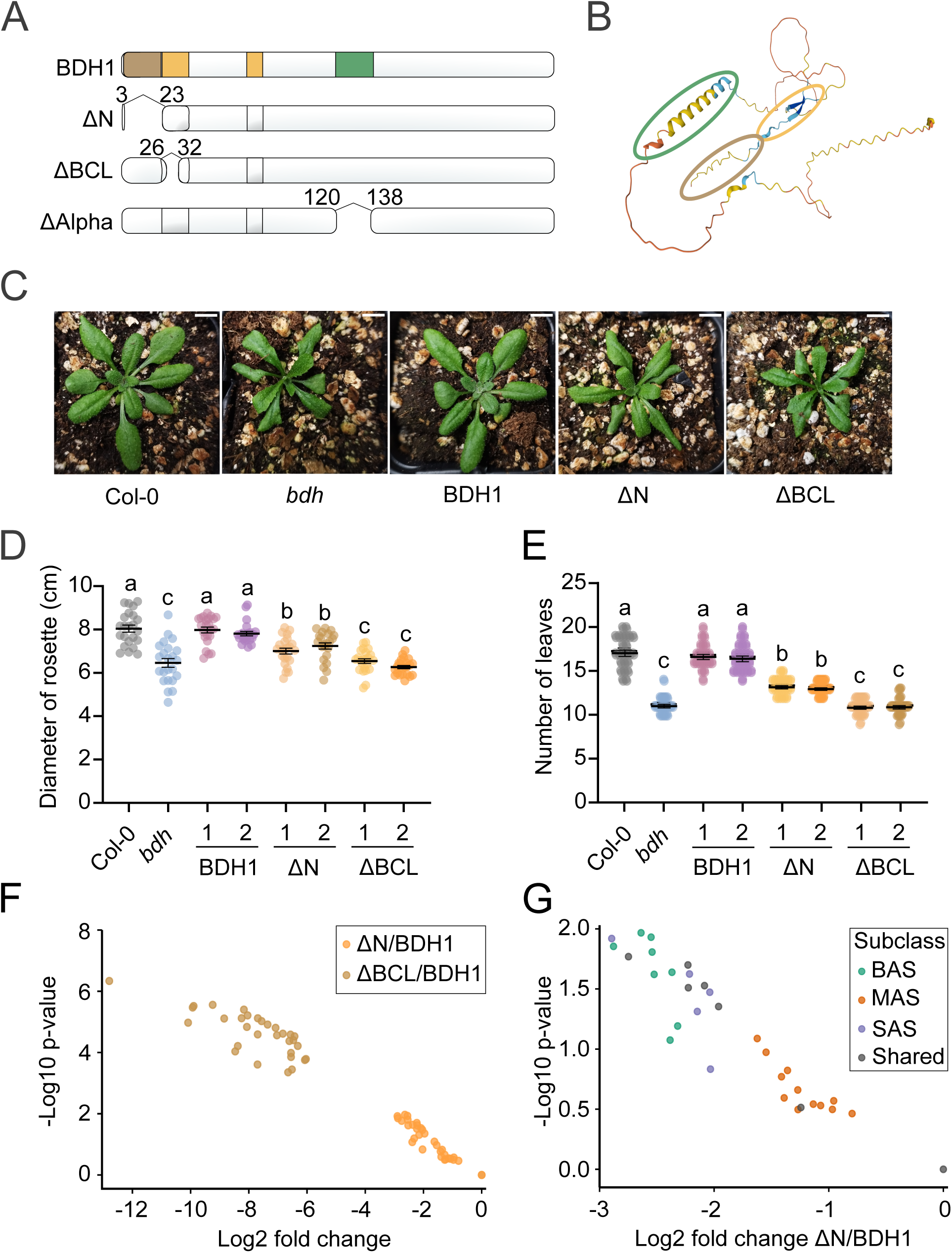
Functional dissection of conserved BDH domains. (A) Schematic representation of the full-length BDH1 protein, depicting the three studied domains: N-domain (brown), BCL-domain (orange), and Alpha domain (green), and the truncated BDH1 versions depicting the first and last aminoacids removed in each deletion. (B) Prediction of the BDH1 protein structure by AlphaFold 2, highlighting the different protein domains following the color scheme described in panel A. Model confidence in the prediction is depicted with different colors: dark blue: very high (pLDDT > 90), light blue: high (pLDDT > 70), orange: low (70 > pLDDT > 50), very low: (pLDDT < 50). (C) Pictures of representative three-week-old plants of the labelled backgrounds: WT, *bdh* double mutant, full length BDH1-3xFLAG (BDH1), N-term-deleted BDH1-3xFLAG (ΔN), and BCL-deleted BDH1-3xFLAG (ΔBCL) (D). Length of the widest measured rosette diameter of the labelled backgrounds in 28-day-old plants grown in long-day conditions. (E) Number of total rosette and caulinar leaves after bolting in the labelled backgrounds. For (D) and (E), the different letters indicate significant differences (P < 0.05), as determined by ANOVA with Tukey’s post-hoc test. Error bars represent Mean ± SEM, n=22-23. Two independent biological replicates were conducted with similar results. (F) Volcano plot depicting the log2 fold change (ΔN/BDH1 and ΔBCL/BDH1) of the intensities of all the SWI/SNF subunits enriched after IP-MS experiments using the ΔN, ΔBCL, and BDH1 transgenic lines. (E) Volcano plot depicting the log2 fold change (ΔN/BDH1) of the intensities of all the SWI/SNF subunits enriched after IP-MS experiments using the ΔN and BDH1 transgenic lines. BAS-, MAS-, and SAS-specific SWI/SNF subunits, as well as those shared in two or three subclasses are depicted in green, orange, purple, and grey, respectively. (F,E) X axis depicts log2 fold change of average intensities of IP experiments. Y axis depicts significance −log10 p-value.

To perform in vivo functional studies of BDH1, we generated a 3xFLAG-tagged BDH1 construct driven by its own promoter and expressed it in *bdh1* single mutants. The resulting transgenic lines were used to profile the BDH1 interactome and genomic targets in inflorescences via IP-MS and ChIP-seq, respectively. Using BDH1-3xFLAG as bait allowed the purification of all subunits from the three described SWI/SNF subclasses, confirming that BDH1 is a common SWI/SNF subunit and suggesting that the generated 3xFLAG-tagged BDH1 transgene was correctly incorporated in the complexes (Supplementary Figure 6A, Supplementary Data 11). Notably, the subunits identified were consistent with those found in a similar experiment performed in seedlings (9, 27), indicating subunit conservation in the three SWI/SNF subclasses across tissues. Furthermore, as expected for a pan-SWI/SNF subunit, ChIP-seq results revealed thousands (14265) of BDH1 targets distributed across the genome in inflorescences (Figure 6B, Supplementary Data 12). A comparison with a previously published BDH1 ChIP-seq in light-grown seedlings (10) (Supplementary Data 13) showed a strong overlap of BDH1 targets, indicating that SWI/SNF remodelers share a large proportion of targets across tissues (Supplementary Figure 6B, Supplementary Data 14). In line with this result, BDH1 co-localized with the three SWI/SNF ATPases in plants -MINU, BRM, SYD-over the same genic region and across thousands of genes, despite these ATPase ChIP-seqs were done in light-grown seedlings(10) (Supplementary Figure 6C).

To investigate the contribution of the BCL, N, and Alpha domains to BDH protein function, we expressed 3xFLAG-tagged full-length BDH1 driven by its own promoter, as well as 3xFLAG-tagged BCL-, N-, and Alpha-domain-deleted BDH1 transgenes (from now on ΔBCL, ΔN, and ΔAlpha, respectively) in *bdh* mutants and investigated their ability to rescue the associated mutant phenotypes. Importantly, the BDH1-3xFLAG genomic fragment was able to rescue all *bdh* mutant phenotypes (Figure 2, Supplementary Figure 7, Supplementary Data 15,16). The deleted BDH1 proteins did not negatively affect protein accumulation and the characterized lines expressed similar or slightly increased BDH1 levels compared to full-length BDH1 (Supplementary Figure 7A, Supplementary Data 15). Notably, ΔAlpha fully rescued defects in silique length in T1 plants and etiolated hypocotyl length in T2 plants to the same extend as full-length BDH1-expressing plants (Supplementary Figure 7B,C, Supplementary Data 15), suggesting no critical function of this domain under the tested conditions. Interestingly, ΔN expression partially rescued these phenotypes, while ΔBCL failed to complement the mutant (Supplementary Figure 7B,C). We then focused on the study of ΔN and ΔBCL as relevant regulators of BDH1 activity. A more comprehensive characterization of these lines in T3 generation, including their ability to rescue other phenotypes like rosette size and flowering time, showed that ΔN and ΔBCL were partially and fully required for BDH1 function (Figure 2C-E, Supplementary Data 16). Further analysis of hypocotyl length in various light conditions (continuous light, LD, SD, and dark) and expression of *XTH* genes in etiolated seedlings revealed that ΔBCL failed to rescue the mutant phenotype or *XTH* expression under any condition, whereas ΔN lines showed varying degrees of complementation being more effective in complementing hypocotyl growth in continuous light and darkness compared to LD and SD (Supplementary Figure 8A-H, Supplementary Data 17).

We hypothesized that the N and BCL domains could be mediating BDH1’s interaction with the SWI/SNF complex. Thus, we performed IP-MS experiments using two independent full-length BDH1, ΔN and ΔBCL transgenic lines. Full-length BDH1 in the *bdh* mutant recovered subunits from all three SWI/SNF subclasses (Supplementary Figure 7D, Supplementary Data 18). Notably, ΔBCL failed to co-purify SWI/SNF subunits, highlighting its requirement for complex interaction and explaining the lack of complementation in ΔBCL lines (Figure 2F, Supplementary Data 18). Conversely, ΔN recovered all three complexes, though with reduced efficiency, indicating an important, albeit not critical, role for the N domain in BDH complex interaction (Figure 2F, Supplementary Data 18). This observation aligns with the partial ability of ΔN plants to complement the *bdh* mutant. Interestingly, the reduced interaction was slightly more pronounced for the BAS and SAS subclasses compared to the MAS subclass (Figure 2G).

### The β-hairpin fold is an evolutionary conserved ACTIN/ARP interacting module in SWI/SNF complexes

Plant BDH and mammal BCL7 proteins exhibit sequence homology within the BCL domain, sharing a similar β-hairpin fold as indicated by their predicted structures (Supplementary Figure 4A,B). However, no additional structural comparisons have been done between these proteins in the context of the complex. Interestingly, yeast SWI/SNF catalytic modules incorporate a subunit known Rtt102 (17), which, despite lacking sequence homology with BDH or BCL7 (12), forms a β-hairpin fold that is reminiscent of the predicted folds of BDH1 and BCL7A BCL domains (Supplementary Figure 9A,B). The crystal structure of Rtt102 in complex with the ATPase (Snf2) HSA domain and the Arp7/Arp9 heterodimer (PDB 4I6M) revealed the molecular mechanism of the interaction between these subunits (17). In the resolved complex, the HSA domain is formed by a long alpha helix which extensively interacts with the ARP heterodimer (Arp7/Arp9) through a central cleft and Rtt102 β-hairpin is positioned in one side of the complex and contacts both ARPs (Figure 3A, Supplementary Figure 9C, Supplementary Data 19). Strikingly, structural modeling of the Arabidopsis and Human SWI/SNF catalytic modules, including BDH1 and BCL7A, respectively, together with their corresponding ACTIN/ARP heterodimers, and ATPase HSA domains (Arabidopsis MINU1 and Human BRG1), resulted in a complex structure resembling the one reported in yeast (Figure 3A-C, Supplementary Figure 9C-E, Supplementary Data 19). The Arabidopsis and Human predicted models showed good prediction values both at the per-residue (pLDDT score) and at the complex level (pTM score) (Supplementary Figure 10A,B). Furthermore, we carried out structural alignments of the experimental yeast complex structure with the predicted AlphaFold2-multimer (31) models, which confirmed very little deviation of the position of the alpha carbons of the models compared to the experimental structure (Supplementary Figure 10C). Like in the yeast complex, the predicted structures show that BDH1 and BCL7 β-hairpins are positioned in a cleft between the ACTIN/ARP heterodimer (Figure 3A-C). The interactions established between these β-hairpins with their respective ACTIN/ARPs are supported by hydrogen bond network and hydrophobic interactions similar to the ones described in the yeast complex (Supplementary Figure 9F-H). Furthermore, supporting their conservation, structural alignments of the predicted BDH1 and BCL7 β-hairpins showed a very strong overlap and relative position of the side chains of conserved key aminoacids (Supplementary Figure 9I). Similarly, BDH1 and Rtt102 alignments predicted a conservation of the overall fold and side chain positions (Supplementary Figure 9J). Notably, there is an equivalent interaction network between β-hairpins and ACTIN/ARPs where conserved tryptophans in the β-hairpins of Rtt102, BDH1, and BCL7, similarly interact with a proline present in one of the two ACTIN/ARP subunits (Arp9, ARP7, and ACTB, respectively) (Figure 3D-F). To experimentally validate these predictions, we performed Co-IP experiments to test the ability of BDH1 to interact with ARP4 and ARP7. As expected, using BDH1 as bait allowed to recover both ARP7 and ARP4 when expressed together (Figure 3G, Supplementary Data 20). Interestingly, BDH1 was able to interact specifically with ARP7 while no interaction was detected with ARP4 when expressed separately (Figure 3H, Supplementary Data 20). This result suggests that the conserved Tryptophan-Proline interaction between the β-hairpins and one of the ARP proteins, in this case between BDH1 and ARP7, might play an important role driving the interaction between BDH/BCL7/Rtt102 and the rest of the complex. Accordingly, this would suggest that BCL7 and Rtt102 preferentially interacts with ACTB and Arp9, respectively (Figure 3E-F). Consistent with the modeled N-term structure of BDH1 when calculated alone (Figure 2B), modeling of the BDH1 N-term region (aminoacids 1-27) in the complex only predicted few hydrogen bond interactions between aminoacids proximal to the β-hairpin (T22, R26) and ARP7 (K347, D351, E352), while the rest of the domain could not be modeled (Supplementary Figure 11A,B). Thus, experimental validation is needed to shed light on the structure and interaction network of the N-term region with other SWI/SNF subunits. This information will be important to understand the impact of the N domain in the ability of BDH1 to interact with the complex. In summary, these results suggest the evolutionary conservation across eukaryotes of the SWI/SNF catalytic module structure and composition.

**Figure 3.**
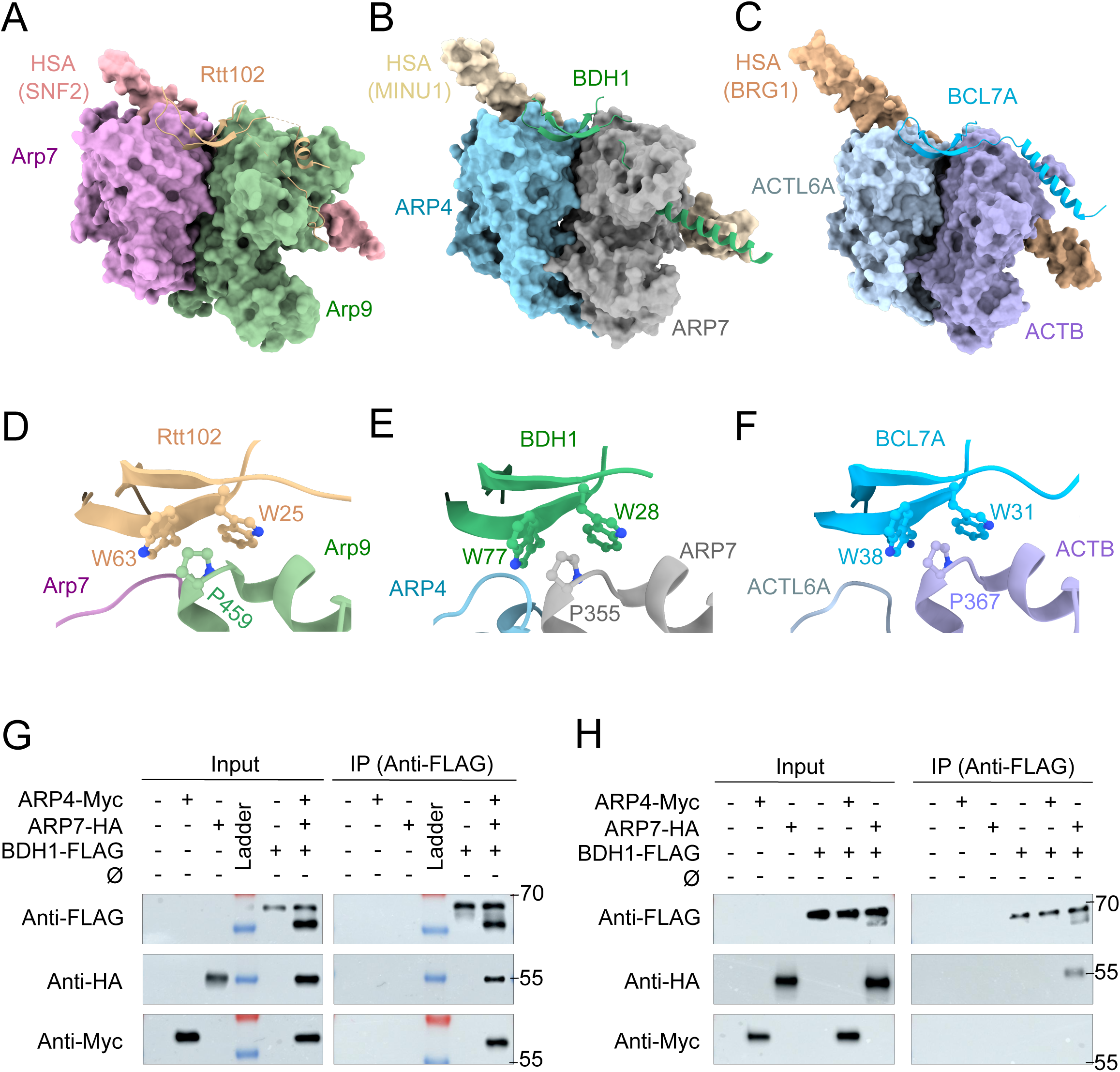
The β-hairpin fold is an evolutionary conserved ACTIN/ARP interacting module in SWI/SNF complexes. (A) Representation of the crystal structure (4I6M) of the *Saccharomyces cerevisiae* SWI/SNF catalytic module depicting Arp7-Arp9-Snf2^(HSA)-^Rtt102. (B-C). Representation of the predicted model of Arabidopsis ARP4-ARP7-MINU1^(HSA)^-BDH1 and Human ACTBL6-ACTB-BRG1^(HSA)^-BCL7A (C) catalytic modules. (D) Detail of the Tryptophan-Proline interaction network observed between Rtt102 and Arp9. Detail of the predicted Tryptophan-Proline interaction between (E) BDH1-ARP7 and (F) BCL7-ACTB. (G) In vivo co-immunoprecipitation (CoIP) of transiently expressed BDH1-FLAG, ARP7-HA, and ARP4-Myc. (H) In vivo CoIP of transiently expressed BDH1-FLAG and ARP7-HA or ARP4-Myc. (G-H) BDH1-FLAG was IP using FLAG antibody and results were observed using anti-FLAG, anti-HA and anti-Myc antibodies. One representative replicate from two independent experiments is depicted.

### Yeast Rtt102 is a distant functional homolog of plant BDH proteins

To further explore the evolutionary conservation between BDH1 and Rtt102 proteins, we investigated whether Rtt102 could functionally replace BDH in plant cells. Structural modeling predicted the Rtt102 β-hairpin would interact with plant ARPs similarly to BDH1-ARP interactions (Supplementary Figure 12A,B). We expressed a 3xFLAG-tagged plant codon-optimized Rtt102 protein (Supplementary Data 21) under the control of the constitutive *UBIQUITIN 10* promoter (*pUBQ10*) in *bdh* mutants and assessed its ability to rescue *bdh* mutant phenotypes. As a control, BDH1 was also expressed under the p*UBQ10* in the *bdh* mutant. Both proteins were properly expressed in T1 plants (Supplementary Figure 12C, Supplementary Data 22). Interestingly, T1 plants expressing Rtt102 slightly rescued some *bdh* phenotypes like the silique defects (Figure 4A, Supplementary Data 23). Remarkably, four independent T2 plants expressing Rtt102 showed an almost complete rescue of the etiolated hypocotyl length defects, similar to full-length BDH1 complementation (Figure 4B, Supplementary Data 23). However, the rescue of other mutant phenotypes, such as rosette size, leaf shape, and flowering time, was significantly reduced (Figure 4C,D, Supplementary Figure 12D, Supplementary Data 22,23). We hypothesized that the weaker complementation by Rtt102 could be due to its reduced affinity for plant ARP proteins. Supporting their functional conservation, Rtt102 specifically interacted with ARP7, although the interaction was weaker than that of BDH1-ARP7, providing a molecular explanation for the partial rescue of *bdh* mutant phenotypes (Figure 4E, Supplementary Data 23). Overall, this data supports the functional conservation between Rtt102 and BDH1 despite their lack of sequence conservation.

**Figure 4.**
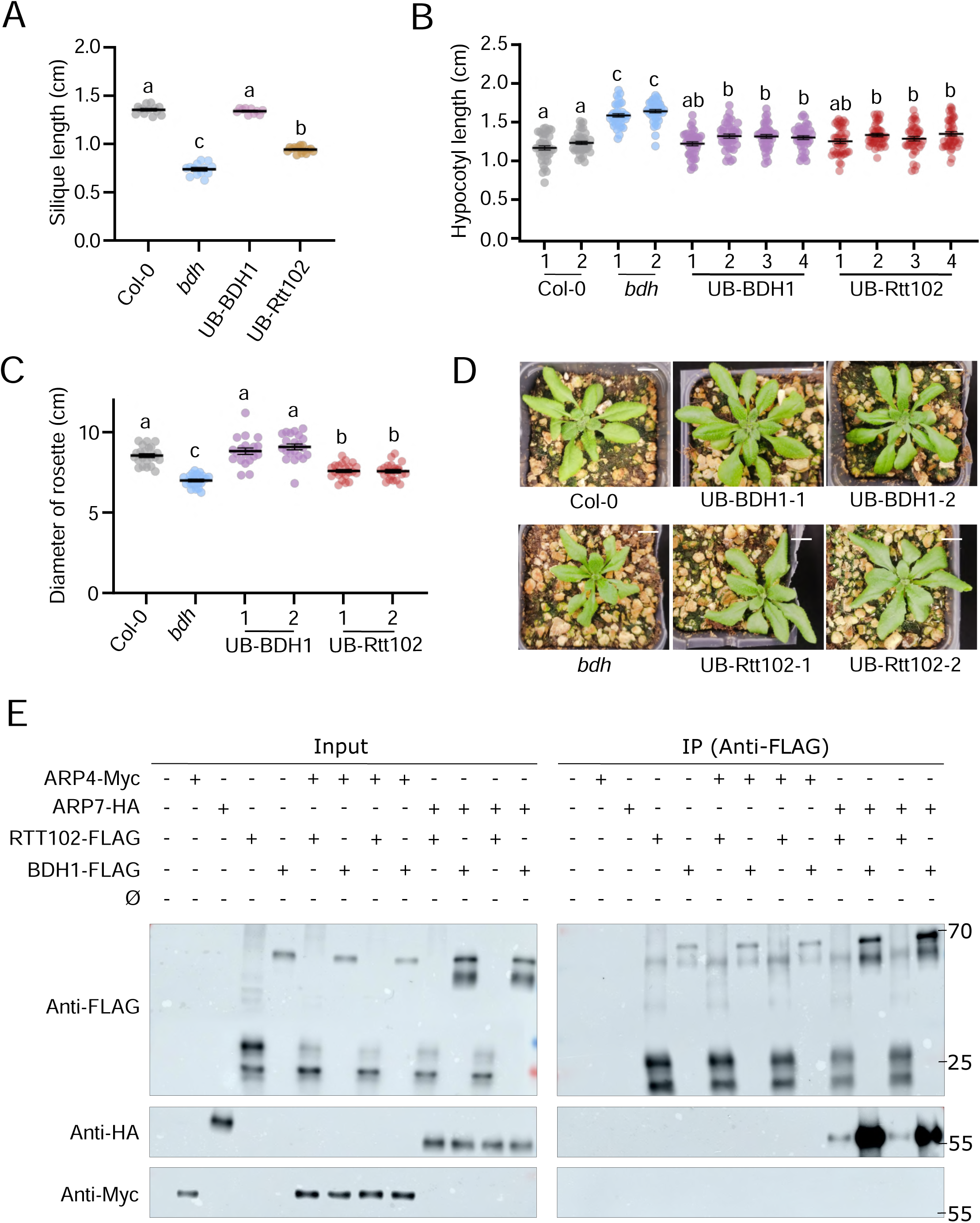
Yeast Rtt102 is a distant functional homolog of plant BDH proteins. (A) Measurement of silique length from the main inflorescence of Col-0, *bdh*, and T1plants expressing UB-BDH1 and UB-Rtt102, n=9-15. (B) Hypocotyl length measurements of 7-day-old etiolated seedlings of Col-0, *bdh*, and four independent T2 populations from UB-BDH1 and UB-Rtt102 lines. (C) Diameter of the rosettes of 28-day-old plants from Col-0, *bdh*, and two independent T2 populations from UB-BDH1 and UB-Rtt102 lines grown under long-day conditions. (D) Top view of representative three-week-old Col-0, *bdh*, and two independent T2 plants expressing pUBQ10::BDH1-6xHA (UB-BDH1) and pUBQ10::Rtt102-3xFLAG (UB-Rtt102). Scale bar: 1 cm. (E) In vivo co-immunoprecipitation (CoIP) of transiently expressed BDH1-FLAG, Rtt102-FLAG, ARP7-HA, and ARP4-Myc. BDH1-FLAG and Rtt102-FLAG were IP using FLAG antibody and results were observed using anti-FLAG, anti-HA and anti-Myc antibodies. Two replicates of the same CoIP experiment are shown.

### BDH promotes ARP heterodimer stability

The predicted position of BDH1 between the two ARP proteins in the catalytic module suggested that it could promote ARP heterodimer interaction. To test this hypothesis, we conducted Co-IP experiments between ARP4 and ARP7 in the presence and absence of BDH1. Notably, the results showed that ARP4 was more efficiently immunoprecipitated by ARP7 when BDH1 was present, suggesting that BDH1 promotes ARP heterodimer formation or stability (Figure 5A, Supplementary Data 24). Next, we investigated whether this effect on ARP heterodimer impacted the overall complex composition. To address this, we generated transgenic plants expressing 6xHA-tagged BAF60B, a core subunit shared by all plant SWI/SNF subclasses and performed IP-MS experiments on these transgenic lines in both WT and *bdh* mutant backgrounds. As anticipated for a plant pan-SWI/SNF subunit, BAF60B isolated subunits from all SWI/SNF complexes (Figure 5B, Supplementary Data 25). Importantly, a significant loss of ARP4 protein was observed when BAF60B was immunoprecipitated in the *bdh* mutant background compared to a similar experiment in WT plants (Figure 5C, Supplementary Data 25). Together with our previous findings, this result supports a model where BDHs promote ARP heterodimer stability, resulting in ARP4 becoming less stable or less incorporated into the complex. This selective loss of ARP4 could explain the molecular and phenotypical defects observed in the *bdh* mutant (9, 26, 27) (Supplementary Figure 1,2).

**Figure 5.**
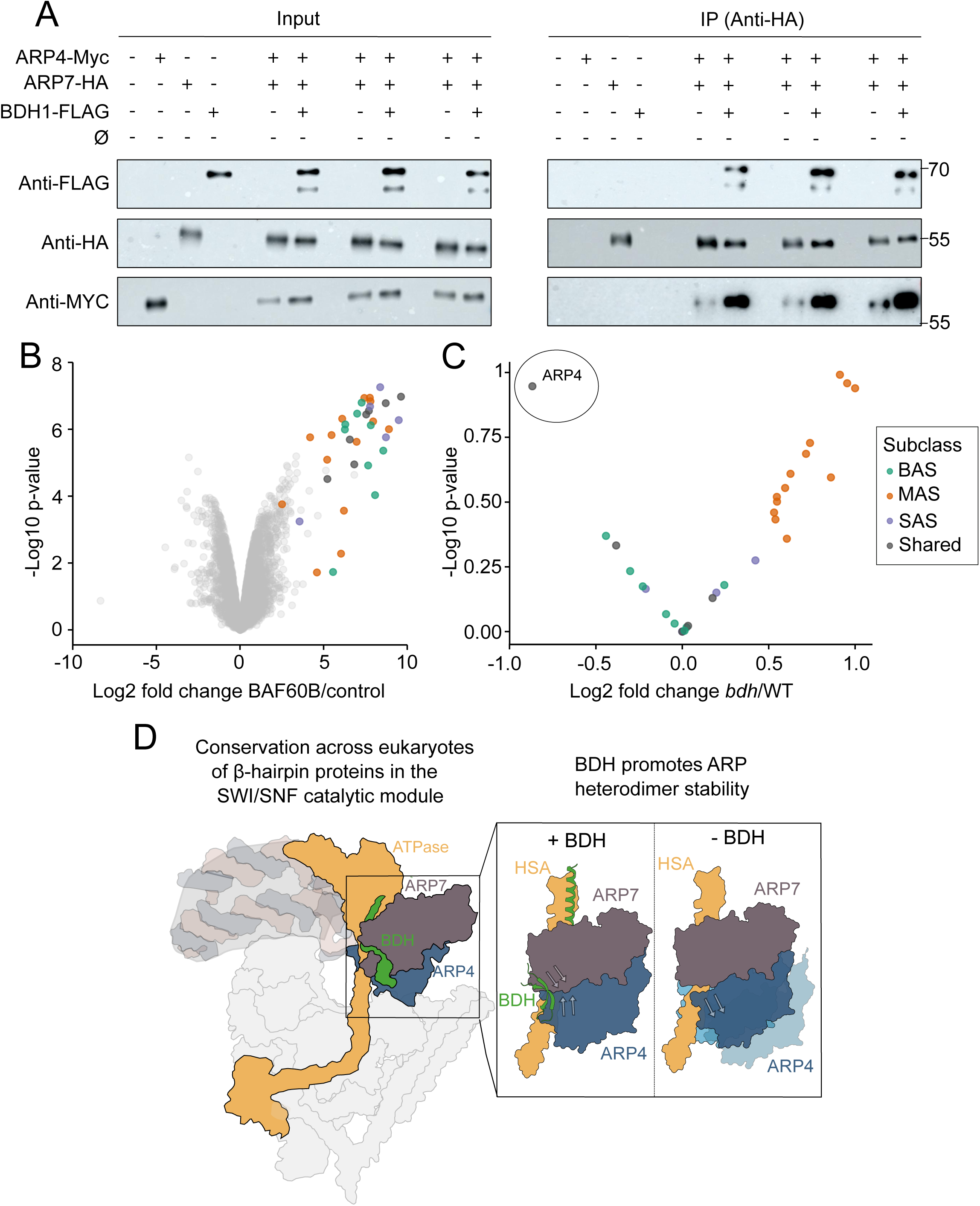
BDH promotes ARP heterodimer stability. (A) In vivo co-immunoprecipitation (CoIP) of transiently expressed BDH1-FLAG, ARP7-HA, and ARP4-Myc. ARP7-HA was IP using HA antibody and results were observed using anti-FLAG, anti-HA and anti-Myc antibodies. Three replicates of the same CoIP experiment are shown. (B) Volcano plot depicting the enrichment (log2 ratio) of all the identified SWI/SNF proteins in the BAF60-6xHA IP-MS experiment in WT background. BAS-, MAS-, and SAS-specific SWI/SNF subunits, as well as those shared in two or three subclasses are depicted in green, orange, purple, and grey, respectively (C) Volcano plot depicting the enrichment of SWI/SNF subunits identified in the IP-MS experiment using BAF60-6xHA as bait in *bdh* mutant vs WT backgrounds. For clarity, ARP4 is enclosed in a circle. X axis depicts log2 fold change of average intensities of IP experiments in WT and *bdh* backgrounds. Y axis depicts significance −log10 p-value. (D) Proposed model for the conservation of β-hairpin proteins in SWI/SNF catalytic modules and their function as promoters of ARPs heterodimer stability. The representation of the complex is based on the CryoEM structure of the BAF mammalian complex65. The position of the ARP heterodimer and BDH protein was inferred from previous models and the prediction reported in our study.

Interestingly, we observed a small increase in the abundance of all the subunits specific to the MAS subclass, whereas the BAS- and SAS-specific subunits showed no or only a slight decrease (Figure 5C). Noteworthy, previous transcriptomic analyses in inflorescences showed that multiple MAS-specific subunits, and not BAS- or SAS-specific subunits, become upregulated in the MAS-specific *tpf* and *minu* mutants (14). Using published RNA-seq data (9), we confirmed this trend in seedlings of *minu* (MAS-specific mutant) and *bdh* mutants (Supplementary Figure 13A) whereas *brm* (BAS-specific) nor *syd* (SAS-specific) mutants did not show such feedback regulation. This indicates that this transcriptional response is uniquely related to MAS subclass malfunction. Thus, the slight increase in MAS subunits accumulation upon BAF60B immunoprecipitation in the *bdh* mutant likely reflects impaired function of the MAS complex as a consequence of BDH depletion.

## Discussion

Decades of research on SWI/SNF chromatin remodelers in animal and fungal models have provided a thorough understanding of the distinct complex subclasses and their subunits (5, 6). Recent studies in plants have confirmed the presence and composition of three plant SWI/SNF subclasses and their equivalence to animal and fungal complexes (9, 10, 12, 14). However, the functions of these plant subclasses and their subunits are only beginning to be understood.

In this work, we characterized the BDH subunit, recently identified as a common component of all three plant subclasses and proposed as a distant ortholog of animal BCL7 proteins (9, 10, 12). We conducted a comprehensive phenotypical characterization to determine the processes affected by the absence of BDH proteins. Additionally, we examined the function of phylogenetically and structurally conserved domains and used modeling to predict the structural conservation of the SWI/SNF catalytic module across eukaryotes. Finally, we investigated the role of BDH proteins in the complex stability, proposing a model in which BDHs promote ARP heterodimer stability.

### BDHs have a broad effect on plant development and participate in hypocotyl cell elongation

We investigated the developmental defects caused by BDH loss of function. Consistent with previous publications (26, 27), we report the redundant function of Arabidopsis BDH1 and BDH2 in controlling multiple phenotypes like the leave shape, flowering time, and silique length. Notably, defects in all these processes were previously described in other plant SWI/SNF mutants but were more severe (9). For example, strong mutants in BRM and SYD ATPases present strong pleiotropic developmental defects that are more pronounced than in *bdh* mutants (32). Furthermore, double *brm* and *syd* mutants or strong *minu1/2* mutants are lethal (32, 33). This suggests that BDH fine-tunes the activities of SWI/SNF subclasses rather than playing a prominent role.

Importantly, we report an overlooked defect in *bdh* mutants, which grow longer hypocotyls. This defect was observed in dark- and light-grown seedlings due to enhanced cell elongation, as hypocotyl cell numbers do not differ between WT and *bdh* mutants. The difference in growth rate between WT and *bdh* mutant occurred after day 4 post-germination, indicating that BDHs facilitate the hypocotyl growth arrest that occurs at this moment of development. In line with these results, light-grown RNAi mutants of the SWI/SNF pan-subunit BAF60B also presented longer hypocotyls, while the opposite was found in BAF60 overexpressors in both light and dark conditions (34). This contrast with the phenotypes found in mutants of different BAS complex subunits (BRM, BRIPs, BRDs) that showed shorter hypocotyls in the dark (35, 36). These results suggest that specific defects in SAS or MAS function, or the combined malfunction of MAS, SAS and/or BAS, could be responsible for the enhanced hypocotyl cell elongation found in *bdh* and *baf60* mutants. Notably, *MINU2* overexpression has been shown to slightly reduce hypocotyl length (37). Transcriptomic analyses revealed that BDH represses the expression of multiple *XTH* genes, which are main players in cell wall shaping and growth. Previous studies showed that overexpression of several Arabidopsis *XTHs* (being, for example, *XTH18* overexpressed just 2-fold compared to WT) was sufficient to promote growth of Arabidopsis etiolated hypocotyls (38, 39). These *XTHs* were among the upregulated genes in our RNA-seq experiment, suggesting that the *XTH* overexpression in the *bdh* mutant could be responsible for the larger hypocotyls. Considering the relevance of XTHs in cell wall shaping and plant growth, understanding how SWI/SNF complexes regulate their expression could have biotechnological applications. While some of these genes are directly bound by BDHs, the mechanistic details about SWI/SNF-mediated regulation of *XTH* expression remain to be explored in future studies. Interestingly, BAF60 was shown to compete with the PIF4 transcription factor for binding to the *XTH15* (aka *XTR7*) promoter to regulate its expression (34).

### Functional characterization of BDH domains

BDH proteins have an evolutionary conserved sequence shared with Human BCL7 proteins, that we named BCL domain (12). Our analyses revealed that the BCL domain is essential for BDH function, likely due to its role in mediating interaction with the ARP heterodimer, particularly with ARP7. Importantly, we only deleted the beta sheet closest to the N-term domain, which includes the conserved W28 shown to mediate interaction with the conserved P355 in ARP7. This result suggests that the formation of an intact β-hairpin fold is required for BDH interaction with the ARPs. Additionally, BDHs have a highly conserved plant-specific region in the proximal N-term region, which is important but not critical for BDH function. Future structural analyses will help revealing how this region folds in the SWI/SNF structure and how it promotes BDH interaction with the complex. Furthermore, BDHs in multiple plant species also have a predicted alpha-helix fold downstream of the β-hairpin region, although this is not conserved at the sequence level. Interestingly, the N-term regions of BCL7 and Rtt102 contain an alpha-helix fold which adopt a similar position within the heterotrimeric complex with ARPs as the one predicted for BDH’s alpha helix, suggesting their functional conservation. However, while N-term deletion of BCL7 fully prevented its function and ability to interact with the complex (40), deletion of the BDH alpha helix had no significant impact on BDH function according to mutant complementation analyses, at least under the conditions/phenotypes tested. This suggests that other regions in BDH, such as its N-term region, might help stabilizing the interaction of BDH with the complex in the absence of the alpha helix domain. Overall, this functional analysis revealed two BDH domains important for its interaction with the complex the complex function.

### Evolutionary conservation of the β-hairpin domain as an ARP-interacting module in SWI/SNF remodelers

The SWI/SNF complexes can be divided into core and catalytic modules (11). While the composition of the core module differs between SWI/SNF subclasses, depending on the incorporation of signature subunits, the catalytic module composition is the same across subclasses, featuring the Snf2-type ATPase and an ACTIN/ARP heterodimer (11, 16, 17). Mammal SWI/SNF complexes incorporate a fourth subunit in this module, BCL7, which has been shown to interact with the ATPase subunit and other complex subunits and the nucleosome (11, 21). Yeast SWI/SNF complexes incorporate Rtt102 in the catalytic module (17), which promotes a more compact ARP heterodimer conformation, enhancing the complex remodeling activity (18, 24). Furthermore, it has been shown to also interact with the ATPase (41). However, despite their similarities, no previous experimental connection had been made between BCL7 and Rtt102 proteins.

A comprehensive phylogenomic analysis across multiple eukaryotic species proposed plant BDH proteins as orthologs of animal BCL7. However, this analysis failed to identify any fungal BDH-BCL7 ortholog (12). In this study, we discovered a structural relationship between animal BCL7, fungal Rtt102, and plant BDH, based on their shared ability to interact with the ACTIN/ARP heterodimer through the formation of a conserved β-hairpin. The link between BDH and Rtt102 is further supported by BDH’s ability to promote ARP heterodimer stability, reminiscent of Rtt102’s function (24). Rtt102 can perform some of the functions of BDH, as demonstrated by *bdh* mutant complementation analyses. Interestingly, the complementation achieved with Rtt102 was similar to that observed with the N-domain-deleted BDH1 (Figure 2, 4, Supplementary Figure 14, Supplementary Data 26). Both were able to rescue the etiolated hypocotyl defects and silique length while only weakly recovering the rosette size and flowering time. Notably, this recovery was to a greater degree than the BCL-deleted BDH1, which completely failed to recover any phenotype (Figure 2, 4, Supplementary Figure 14, Supplementary Data 26). This functional connection between Rtt102 and N-domain-deleted BDH1 can be attributed to their weaker ability to interact with the complex compared with full-length BDH1 (Figure 4A, Figure 2F,G). The phenotype-dependent ability of Rtt102 or N-domain-deleted BDH1 to rescue the *bdh* mutant might reflect distinct modes of action or catalytic activities of the SWI/SNF complexes that may rely on BDH1 function to varying degrees, potentially acting on specific processes or genomic contexts.

Interestingly, a recent study reported the structural conservation of a similar ARP-β-hairpin interacting module in the context of the animal and fungal INO80 complexes (42). This study showed that the animal and yeast β-hairpin-containing proteins YY1 and Ies4, respectively, interact through two conserved tryptophans with a conserved proline in one of the two ARPs that heterodimerize in the animal and yeast complexes (ACTB-YY1 and Arp4-Ies4). Notably, our study shows that BCL7, Rtt102, and BDH also conserve these tryptophans which are predicted to interact specifically with one of the two ARPs in a similar manner. Notably, to our knowledge, no functional tests are available on the role of YY1 or Ies4 in the Actin/ARP heterodimer stability. Similarly, the Snf2^HSA^-ARP heterodimer arrangement is also conserved in the SWR1 and NuA4 complexes, both of which incorporate the β-hairpin containing protein Swc4 (43, 44). What gives specificity to the incorporation of these β-hairpin proteins to each complex could be dictated by their stronger affinity of the β-hairpin protein to the complex-specific ARP protein. For example, BDH1 protein strongly interacts with the ARP subunit that is specific to the SWI/SNF complex, ARP7, and not with ARP4, also found in INO80, SWR1 and NuA4 complexes (43, 44). Similarly, Rtt102 showed a stronger interaction with the yeast SWI/SNF-specific ARP, Arp9 (24, 45). According to our modeling, this logic would predict that BLC7 interacts more strongly with ACTB. An alternative situation occurs in fungi where INO80 and SWI/SNF complexes incorporate the same actin-Arp4 heterodimer. In this case, a specific region in Ies4 was shown to provide complex specificity through the interaction with an additional INO80-specific subunit, ARP8, which is found near the actin-Arp4 module (42).

### BDH promote stability of the ARP heterodimer in the catalytic module

A recent study reported genome-wide changes in chromatin accessibility and gene-body nucleosome occupancy in Arabidopsis *bdh* mutants (27). However, this study did not find substantial changes in the BAS complex composition or its recruitment to the chromatin in the absence of BDH, thus leaving unanswered how BDH depletion impaired SWI/SNF function. We propose that BDH impacts SWI/SNF function through the stabilization of the ARP heterodimer. We showed that ARP7-ARP4 interaction is enhanced in the presence of BDH1 and that BDH depletion triggers ARP4 destabilization in vivo. The partial loss of ARP4 observed in IP-MS experiments could reflect an overall weaker ARP heterodimer stability across all SWI/SNF subclasses. Notably, experiments in yeast showed that Rtt102 promotes a more compact ARP conformation which, in turn, shortens the interaction surface of the heterodimer with the HSA domain, altering the network of interactions between distinct ATPase domains important for catalytic activity, like post-HSA and Protrusion 1 (24). Importantly, plant BDH, as well as BCL7 and Rtt102, can also interact with the ATPase which could also directly influence its activity (21, 27, 41). Future mechanistic studies will shed light on the specific impact of BDH on ARP compaction, ATP turnover, DNA translocation, and nucleosome remodeling in plants. Additionally, BCL7 proteins have been shown to interact with other proteins in the complex and the nucleosome, such as SMARCB1, H2B and H2A (11, 21). Although we do not yet know whether BDHs interact with all these proteins, it is plausible that BDH depletion leads to altered subunit contacts within the complex and with the nucleosome, resulting in remodeling defects.

Apart from the decrease in ARP4, we observed a slight increase in MAS-specific subunits isolated by BAF60B in the *bdh* mutant compared to WT. This might reflect a small increase in the accumulation of the MAS complex in the *bdh* mutant. Importantly, malfunction of the MAS complex leads to upregulation of MAS-specific subunits but not BAS- and SAS-specific ones (14) (Supplementary Figure 13A). Thus, these results suggests that, whereas a small increase in MAS complex might occur in *bdh* mutants, these complexes are functionally impaired, leading to the transcriptional upregulation of MAS-specific subunits. On the contrary, we did not observe significant changes between WT and *bdh* in the accumulation of BAF60B-immunoprecipitated BAS- and SAS-specific subunits. In line with this result, a recent study found no differences in BAS-specific subunit accumulation when pulling from BRM in WT and *bdh* mutant (27). Interestingly, these IP-MS results showed a specific reduction in ARP4 peptides compared to other BAS subunits identified, supporting our observation that BDH alters ARP4 stability in the complex. Furthermore, this study showed that BDH affected chromatin accessibility in targets of the three subclasses (27). A clustering analysis of RNA-seq experiments conducted on multiple plant SWI/SNF mutants (9) revealed that DEGs in *bdh* mutants exhibit expression changes more closely aligned with BAS complex mutants (*brm, an3, brd1/2/13, brip1/2*) followed by SAS (*syd, swi3d, sys*) and then MAS complex mutants (*minu1/2, tpf1/2*) (Supplementary Figure 13B). This indicates that, although BDH influences MAS complex function, leading to the upregulation of MAS subunits in its absence, it may have a stronger impact on BAS and SAS function.

In summary, our findings suggest that BDHs influence the catalytic activity of plant SWI/SNF subclasses by stabilizing the pan-SWI/SNF ARP heterodimer module (Figure 5D). Given the analogous role of yeast Rtt102 in Arp heterodimer compaction (24) and the structural conservation indicated by our study, we propose that BCL7, BDH, and Rtt102 are functionally equivalent subunits of the SWI/SNF catalytic modules conserved across eukaryotes.

## Material and Methods

### Plant materials and Growth conditions

The *bdh1-1* (SALK_152173) and *bdh2-1* (SALK_029285) mutants used in this study are in the Col-0 background, were previously characterized (26), and were requested from ABRC. A double *bdh1/bdh2*, called *bdh* in this manuscript, was obtained by genetic cross. Oligos for T-DNA genotyping are found in Supplementary Data 27. Plants were grown in a growth chamber set to a 16L/8D light cycle (LD) and 22°C. For in vitro growth, seeds were sterilized with chlorine gas and were plated on 0.5x Murashige and Skoog medium (MS, Caisson Laboratories) containing 0.8% agar (Sigma) adjusted to pH 5.7, followed by incubation at 4°C in darkness for 3 days. After stratification, plates were moved to a Percival incubator (Percival-scientific.com) at 22 °C with different light cycles, qualities, and intensities. Light intensities were measured with a Spectrosense2 meter (Skye Instruments Ltd). Transgenic lines were selected on 0.5x MS medium supplemented with BASTA (6.67 μg/ml) (Sigma) or Hygromycin (35 μg/ml) (Applichem).

To assess hypocotyl length, seedlings were grown evenly spaced on plates under the specified light conditions and photoperiod. After the designated period, the seedlings were scanned and the resulting images were analyzed using the ImageJ software (https://imagej.nih.gov.ij). The number of hypocotyl cell was counted with a Nikon Eclipse Ni microscope. To measure flowering time, total number of rosette and caulinar leaves were counted after bolting. To measure rosette diameter, images of 28-day-old plants were taken and the diameter was counted as the length from tip to tip of the longest rosette leaves.

### Construction of plasmids and transgenic lines

The BDH1 genomic region, including 2030 bp upstream of the start codon and all the gene body, including introns, until the stop codon, was amplified from genomic DNA of Col-0 plants and cloned into a pENTR/D vector (Invitrogen) to generate pENTR-gBDH1. The primers used for this and the following clonings can be found in Supplementary Data 28. Subsequently, the BDH1 fragment was transferred via LR reaction into a modified pEarleyGate302 vector containing a 3xFLAG tag downstream of the gateway cassette (pEG302-3xFLAG) to produce pEG-gBDH1-3xFLAG. This construct was then transformed into *Agrobacterium tumefaciens* strain GV3101 C58C1 to transform the *bdh1-1* mutant employing the floral dip method. To construct the Δalpha, ΔBCL, and ΔN plasmids, primers were designed to re-amplify the pENTR-gBDH1 plasmid excluding the regions indicated in Figure 2B, and the resulting fragment was then re-ligated with T4 DNA ligase (Thermo Scientific). The resulting constructs were transferred into pEG302-3xFLAG using LR reaction to generate pEG-gBDH1ΔAlpha-3xFLAG, pEG-gBDH1ΔBCL-3xFLAG, and pEG-gBDH1ΔN-3xFLAG constructs. Subsequently, these constructs, along with pEG-gBDH1-3xFLAG, were introduced into *Agrobacterium tumefaciens* strain GV3101 C58C1 to transform the *bdh* double mutant.

Gateway-compatible vectors that incorporate the SWI/SNF subunits were requested from ABRC as bacterial stabs (*ARP4* (G23361), *ARP7* (U24661), *BAF60B* (G15375), *BDH1* (G50477),). The stop codon present in the CDSs of pENTR-ARP7, pENTR-BAF60B, and pENTR-BDH1 was removed by site-directed mutagenesis. The resulting plasmids were transferred by LR reaction (Invitrogen) into the pB7m34GW or pH7m34GW binary destination (https://gateway.psb.ugent.be) together with a *UBQ10* promoter fragment in the 5’position and a 6xHA tag in the 3’position to create pB7-UBQ10::ARP7-6xHA, pH7-UBQ10::BAF60B-6xHA, and pB7-UBQ10::BDH1-6xHA. pENTR-ARP4 was transferred to pEarleyGate203 (46) by LR reaction to create pEG203-ARP4. A pENTR containing a plant codon-optimized Rtt102 fused to 3xFLAG sequence in its C-terminus was synthesized from Genescript (Sequence of the codon optimized Rtt102-3xFLAG can be found in Supplementary Data 21). The Rtt102-3xFLAG fragment was transferred to the pB7m24GW binary destination (https://gateway.psb.ugent.be) together with a *UBQ10* promoter fragment in the 5’position to create pB7-UBQ10::Rtt102-3xFLAG.

### Transient expression in Nicotiana benthamiana

*Agrobacterium tumefaciens* GV3101 C58C1 cells harboring the relevant constructs were cultured overnight at 28°C in liquid LB medium supplemented with appropriate antibiotics. Following pelleting, the cultures were resuspended in 10 mM MES-KOH buffer at pH 5.6, containing 10 mM MgCl2 and 150 µM acetosyringone, to achieve a final OD600 of 0.5, followed by a 2 h incubation at room temperature in the dark. Subsequently, the relevant suspensions were mixed and infiltrated into *Nicotiana benthamiana* leaves at a final OD600 of 0.1 each. All experiments were co-infiltrated with an Agrobacterium strain transformed with a plasmid to express the p19 silencing suppressor at a final OD600 of 0.05. Samples were collected 3 days post-inoculation.

### Western blot detection

Ground samples in liquid nitrogen were used to extract total proteins by resuspending the tissue in SDS-PAGE buffer followed by denaturation at 95°C for 5 minutes. Samples were resolved on 4–10% SDS-PAGE gels. Subsequently, proteins were transferred onto PVDF membranes (Amersham) and stained with Ponceau S to confirm efficient transfer and assess loading consistency. The membranes were then probed with either Horse Radish Peroxidase (HRP)-conjugated 3F10 anti-HA (1:2000, Roche), HRP-conjugated FLAG M2 (1:5000, Sigma) or Anti-Myc-HRP 9E10 (1:5000, Invitrogen). Chemiluminescent signals were developed using Supersignal Dura and Atto substrates (Thermo Scientific) and visualized with an ImageQuant 800 system (Amersham).

### Protein co-immunoprecipitation

Agroinfiltrated Nicotiana leaves were ground to a fine powder in liquid nitrogen using a mortar and pestle, and then resuspended in 1.5 ml of IP buffer (50 mM Tris-HCl pH 7.5, 150 mM NaCl, 5 mM MgCl2, 0.1% NP-40, 10% glycerol, 0.5 mM DTT, 1 mM PMSF, 1x Complete Mini EDTA-Free Protease Inhibitor (Roche). The extracts were clarified twice by centrifugation at 12000 × g for 5 min at 4°C. Total protein concentration was determined using the Bradford Protein Assay (Bio-Rad) and normalized to 0.750 mg/ml. An amount between 0.5-1.5% of the extracts were set aside for protein level verification (input). For immunoprecipitations, the extracts were incubated with 1 µg of anti-FLAG M2 monoclonal antibody (Sigma) or anti-HA 3F10 monoclonal antibody (Roche) for 2 h with gentle rotation at 4°C. Subsequently, 20 µl of pre-washed magnetic protein G Dynabeads (Invitrogen) in IP buffer were added to the samples and incubated for 1 h with gentle rotation at 4°C. The samples were then washed three times with IP buffer, and the precipitated proteins were eluted by heating the beads at 95°C for 2 min in 50 µl of 2x SDS-PAGE loading buffer. Finally, 5 and 20 µl of the eluate were separately subjected to Western blot analysis to detect the immunoprecipitated and co-immunoprecipitated proteins, respectively.

### Immunoprecipitation and mass spectrometry (IP-MS)

Immunoprecipitation coupled with mass spectrometry was conducted following established protocols (12, 14). Briefly, 7 grams of inflorescences were ground in liquid nitrogen and resuspended in 40 ml of IP buffer (50 mM Tris pH 7.5, 150 mM NaCl, 5 mM MgCl2, 10% glycerol, 0.1% NP40, 0.5 mM DTT, 1 mM PMSF, 1 μg/μl pepstatin, and 1× Complete EDTA-Free Protease Inhibitors (Roche). Samples were filtered using one layer of Miracloth (Merck, cat#475855), homogenized with a douncer (10 times soft, 10 times hard), and then centrifuged at 4°C for 10 min at 10,000 × g. The supernatant was filtered again using a 40 μm cell strainer. Next, 200 µl of Anti-FLAG M2 magnetic beads (Sigma, cat#M8823) or 250 µl of Pierce Anti-HA magnetic beads (Thermo Scientific), to immunoprecipitated BDH1-3xFLAG (and BDH1 deletions) or BAF60B-6xHA, respectively, were pre-blocked with 5% BSA, and were added to the samples, which were rotated at 4°C for 3 h. The samples were washed four times with IP buffer and two times with IP buffer without NP40. Elution of BDH1-3xFLAG associated complexes was performed with 300 μl of 250 μg/ml 3xFLAG peptide (Sigma, cat#F4799) in IP buffer without NP40 for 30 min at 25°C, repeated once with a 15 min incubation at 37°C. TCA was added to a final concentration of 20%, followed by a 30 min incubation on ice and a subsequent 30 min centrifugation at 4°C at 12,000 × g. Samples were then washed three times with 250 μl of cold acetone, and the pellet was air-dried. Elution of BAF60-6xHA was directly done in 200 μl of 1x SDS-PAGE buffer followed by incubation at 95°C for 10 min.

### Proteomic characterization of affinity purified protein complexes

BDH1-3xFLAG (and BDH1 deletions) and BAF60-6xHA Immunoprecipitates were processed in two ways. BDH1 (and deletions) immunoprecipitates were processed essentially as previously describe (47). Acetone precipitated immunoprecipitates were resuspended in 8M urea, 100 mM Tris-HCl, pH 8.0 buffer, diluted to 2M urea with 100 mM Tris-HCl, and then reduced and alkylated by the sequential addition of 5 mM tris(2-carboxyethyl)phosphine and 10 mM iodoacetamide. Samples were then digested overnight at 37°C with lys-c and trypsin. Digested peptides were then desalted using peptide-level SP3 and analyzed by LC-MS/MS on a Orbitrap Fusion Lumos mass spectrometer (48). BAF60-6xHA samples were eluted in SDS Laemmli buffer, diluted 1:2 with 100mM Tris-Cl, pH 8.0, reduced and alkylated, and then isolated using protein-level SP3. The samples were then digested on-bead with lys-C and trypsin proteases overnight at 37°C followed by peptide desalting using SP3. BAF60-6xHA immunoprecipitates were analyzed on a Bruker timsTOF HT mass spectrometer.

For LC-MS/MS analysis of BDH1 (and deletions) on the Orbitrap Lumos, peptides were fractionated on a C18 reversed phase fused silica capillary column (75 μm inner diameter) packed in-house to a length of 25 cm with bulk 1.9mM ReproSil-Pur beads with 120 Å pores (49). Peptides were eluted with an acetonitrile gradient generated using a Dionex Ultimate 3000 (Thermo Scientific) HPLC at a flow rate of 200nL/min. The MS/MS spectra were collected using data dependent acquisition with a MS1 (R=120,000) scan to identify precursors of interest followed by sequential MS2 (R=15,000). MS/MS data were processed using MSFragger integrated in the Fragpipe computational environment as described previously (50). Data was searched using *Arabidopsis thaliana* database (Uniprot Reference UP000006548) and filtered using peptide and protein level false discovery rates of 0.01. Quantification was performed with ionquant using MS1-based LFQ (51). Fold-change, normalization, and significance testing calculations were performed using the combined_protein.tsv output obtained from MSFragger and the Fragpipe-analyst package (52).

Analysis of the BAF60-6xHA IPs was performed on a Bruker timsTOF HT using diaPASEF (53). Peptides were separated using ionopticks AuroraElite C18 Columns (75µm ID, 15 cm lengths, 1.5 μm particle size) using a Vanquish Neo UHPLC-System (Thermo Scientific) running at 500nL/min. Data was acquired on a Bruker timsTOF HT mass spectrometer using a diaPASEF data acquisition strategy. dia-PASEF windows were 10m/z wide and spanned the range of 300 m/z to 1200 m/z. Ion mobility parameters were as follows: 1/K0 start: 0.60 Vs cm−2; 1/K0 end: 1.60 Vs cm−2; ramp time: 100 ms; accumulation time: 50 ms. Database searching was done using DIA-NN and an *in silico* library generated from a *Arabidopsis thaliana* database (Uniprot Reference UP000006548) (54). Fragpipe-analyst was used to identify differentially enriched proteins from DIA-NN output files (52).

### RNA extraction, qRT-PCR and RNA-seq libraries

Total RNA was isolated using the Direct-zol RNA Miniprep Kit (Zymo Research) according to the manufacturer’s instructions. For *XTH* expression analyses, 1 μg of total RNA from 5-day-old etiolated seedlings of the indicated backgrounds was used for cDNA synthesis with the NZY First-Strand cDNA Synthesis kit (NZYTech) following the manufacturer’s protocol. Primer details are provided in Supplementary Data 27. Fold change was determined relative to the expression of the *PP2A* housekeeping gene using the ΔΔCt method (55). For RNA-seq experiments, total RNA from 5-day-old etiolated seedlings of Col-0 and *bdh* backgrounds (three biological replicates per background) was submitted to the BGI company for the preparation of strand-specific mRNA libraries, which were sequenced using the DNBSEQ high-throughput platform as PE100 reads.

### RNA-seq data analysis

For RNA-seq data analyses, reads were mapped to the TAIR10 genome using STAR (56) v2.7.10b with default options (Supplementary Data 28). Read counts were obtained with the featureCounts (57) command with options “-Q 5” and “-p --countReadPairs” in the case of paired-end data and using the Araport11 annotation. Counts were normalized with the ‘rlog’ function from DESeq2 (58) v1.38.0. A Principal Component Analysis (PCA) was performed over these counts using the ‘plotPCA’ function from DESeq2 (58) v1.38.0. Differential expression analyses were performed using DESeq2 1.38.0 with default options. Fold changes were corrected using the lfcShrink function with the ‘apeglm’ method (59). GO enrichment analyses over DEGs were performed using the ShinyGO (60) v0.80 web tool, selecting GO Biological Process as database and default parameters.

### Chromatin immunoprecipitation

The chromatin immunoprecipitation (ChIP) protocol was conducted according to previously described methods with minor adjustments (14). Briefly, 1 gr of inflorescences of Col-0 and two independent BDH1-3xFLAG transgenic lines were ground in liquid nitrogen and crosslinked in Nuclei Isolation Buffer containing 1% formaldehyde for 10 min at room temperature. The crosslinking reaction was quenched with glycine, followed by nuclei isolation and chromatin shearing using a Bioruptor Pico (Diagenode). Chromatin was then immunoprecipitated overnight at 4°C with Anti-FLAG M2 antibody (5 μl/ChIP, F1804, Sigma). The immunocomplexes were captured using a 1:1 mixture of magnetic Protein A and Protein G Dynabeads (Invitrogen) for 3 h at 4°C, followed by sequential washing with low salt, high salt, LiCl, and TE buffers for 10 min each at 4°C. Elution of the immunoprecipitated complexes was performed for 2 × 20 min at 65°C with elution buffer. Reverse crosslinking was carried out overnight at 65°C, followed by proteinase K treatment at 45°C for 5 h. DNA was purified using phenol:chloroform:isoamyl alcohol 25:24:1 (Fisher Scientific) and precipitated with GlycoBlue (Invitrogen) and NaAc/EtOH overnight at −20°C.

Samples were spun for 30 min at max speed at 4°C followed by a 70% ethanol wash. The purified DNA was resuspended in 75 μl of elution buffer. Libraries for sequencing were prepared using the Ovation Ultra Low System V2 1–16 kit (NuGEN) following the manufacturer’s instructions. Libraries were sequenced in a HiSeq 2500 as SE50 reads.

### ChIP-seq data analysis

For ChIP-seq data analyses, reads were aligned to the TAIR10 genome using bowtie2 (61) v2.5.2 with default parameters (Supplementary Data 28). Duplicates were subsequently marked using the sambamba markdup command from sambamba v1.0 (62). Then, uniquely mapped reads were retained by filtering the alignments with the samtools view command from samtools (63) v 1.17 with options “-F4 -F 1024 -q 5”. Coverage tracks of these alignments were generated using the bedtools genomecov command from bedtools (64) 2.31.0 with parameters “-bga -fs 200”. The coverage tracks were normalized to Counts Per Million (CPM). Correlation between ChIP-seq replicates was estimated by first counting the reads over 500 bp windows with the multiBigwigSummary command from deeptools and then calculating the Pearson correlation coefficient (r) between them with the cor() function in R. The signal of ChIP-seq replicates showed a high correlation genome-wide (r = 0.95). Biological replicates were averaged using the wiggletools mean command from wiggletools (65) 1.2.11. The BedGraph files were converted to bigwig with the bedGraphToBigWig utility from UCSC tools. Peak calling was performed on clean alignments using macs2 (66) v2.9.1 with parameters “--nomodel --extsize 200”. Peaks from biological replicates were merged with the bedtools merge command, and the consensus peaks (present in both replicates) were annotated to their closest gene using the bedtools closest command. Genes that intersected with these peaks or were less than 2 kb away from peak summits were considered as targets.

### Structural modeling and analysis

The Arabidopsis (MINU1^HSA^-ARP4-ARP7-BDH1) and Human (BRG1^HSA^-ACTBL6-ACTB-BCL7A) complexes were modelled with AlphaFold2 (29) and AlphaFold2-multimer (31) using a colab notebook running ColabFold (67) v1.5.5. For model quality assessment, the predicted local distance difference test (pLDDT) and the predicted aligned error were calculated for each model (Supplementary Figure 10A,B). Structural alignment of the predictions against the experimental structure of yeast (Snf2^HSA^-Arp7-Arp9-Rtt102) (PDB 4I6M) was carried out in ChimeraX (CITE) to further asses the models (Supplementary Figure 10C). The root mean square deviation (RMSD) of the Cα (in Angstroms) between the models and the yeast crystal structure PDB 4I6M was calculated in ChimeraX. The amino acid sequences used for complex prediction are provided in Supplementary Data 19.

### Statistical analyses

All statistical analyses conducted in this manuscript are described above. Methods for statistical tests, sample sizes, p-values are provided in the figures. A two-tailed Student’s t-test was used to evaluate the differences between the two groups. For multiple comparisons, a one-way analysis of variance (ANOVA) was conducted, followed by Tukey’s honestly significant difference (HSD) post hoc test. Statistical analyses were conducted using GraphPad Prism software (v.8) and Rstudio. All experiments were conducted with a minimum of two replicates.

### Figure representation

Venn diagrams were drawn using eulerr (https://eulerr.co/). Heatmaps and metaplots were produced using the computeMatrix and plotHeatmap or plotProfile commands from deeptools (68) v3.5.1. Plots of RNA-seq data and IP-MS volcano plots were drawn using ggplot2. The protein alignments were performed using CLC Main Workbench 24 with a gap open cost of 10.0 and a gap extension cost of 1.0.

## Data availability

The ChIP-seq and RNA-seq have been deposited at GEO with accession numbers GSE268510 and GSE268511, respectively. The ChIP-seq data of BDH1, MINU2, BRM and SYD in seedlings was obtained from GSE218841. The RNA-seq data of *bdh* and WT seedlings was obtained from GSE193095. The IP-MS data have been deposited at Massive Arabidopsis.

## Author contributions

J.C-F performed all experiments. J.P.-A. performed all bioinformatics analyses. B.D-M participated in expression analyses. J.L-J performed structural modeling and analyses. V.P. and J.W. performed the IP-MS analyses. J.C-F. and J.G.-B. conceived this study and wrote the manuscript.

