## Supplementary Figures for "Plant BCL-Domain Homologues play a conserved role in SWI/SNF complex stability"

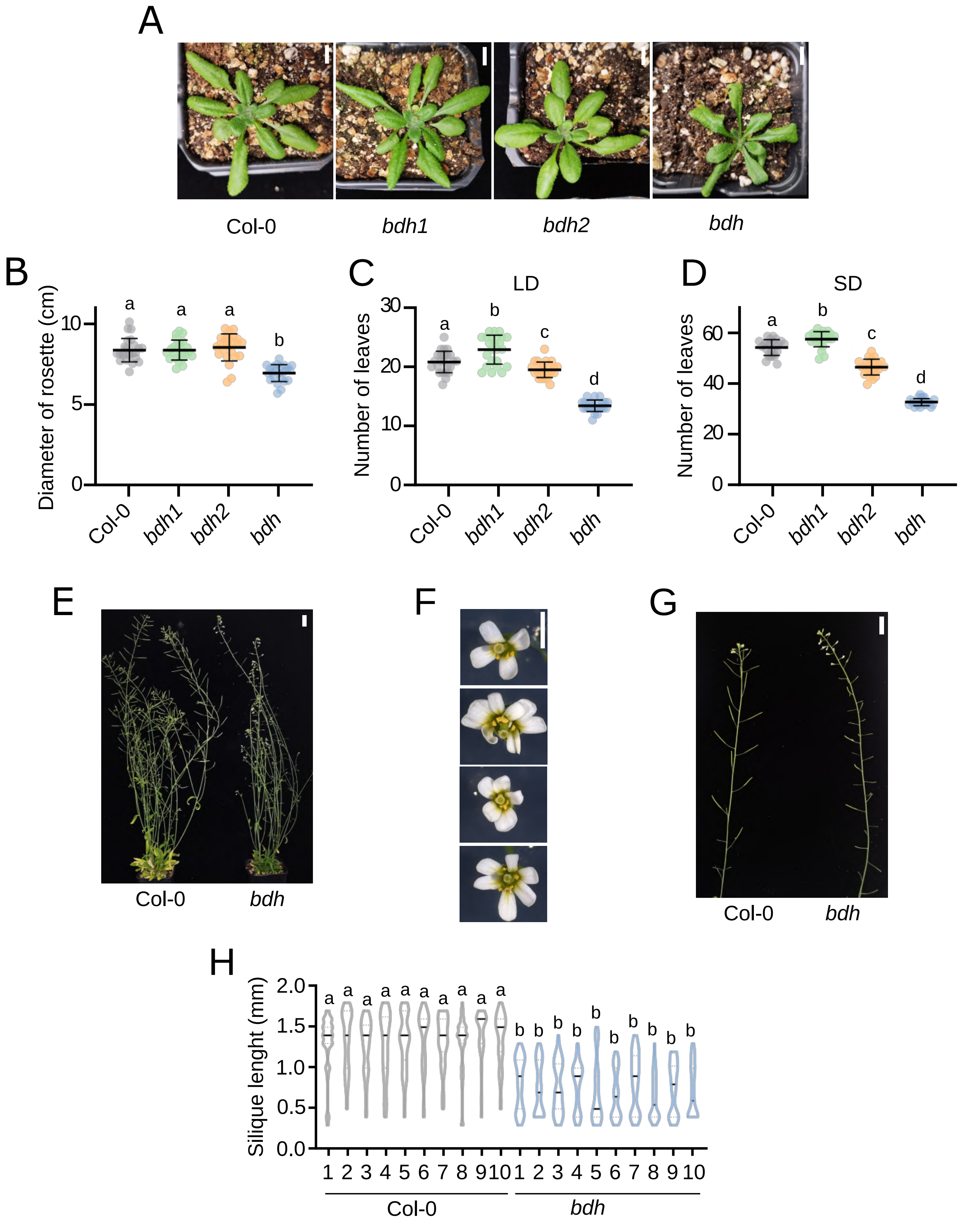


**Supplementary Figure 1. BDH regulate multiple developmental processes**

(A) Top view of 25-day-old *Col-0, bdh1, bdh2 and bdh (bdh1/2)* mutants, scale bar: 1 cm (B) Diameter of the rosettes of the labelled backgrounds grown under long-day conditions. (C,D) Flowering time reported as number of leaves (rosette+caulinar) after bolting under long-day (C) and short-day conditions (D) in the labelled backgrounds. (E) 7-week-old representative Col-0 and *bdh* adult plant grown in soil under long-day condition, scale bar of 2 cm. (F) Representative images of *bdh* mutant flowers depicting a normal number of petals (upper panel) and extra petals. Scale bar of 0.5 mm. (G) Representative primary inflorescences of Col-0 and *bdh* plants. Scale bar of 2 cm. (H) Measurement of silique length in the primary inflorescence of Col-0 and *bdh* mutants, n=10. Student's t-test, *P < 0.05.

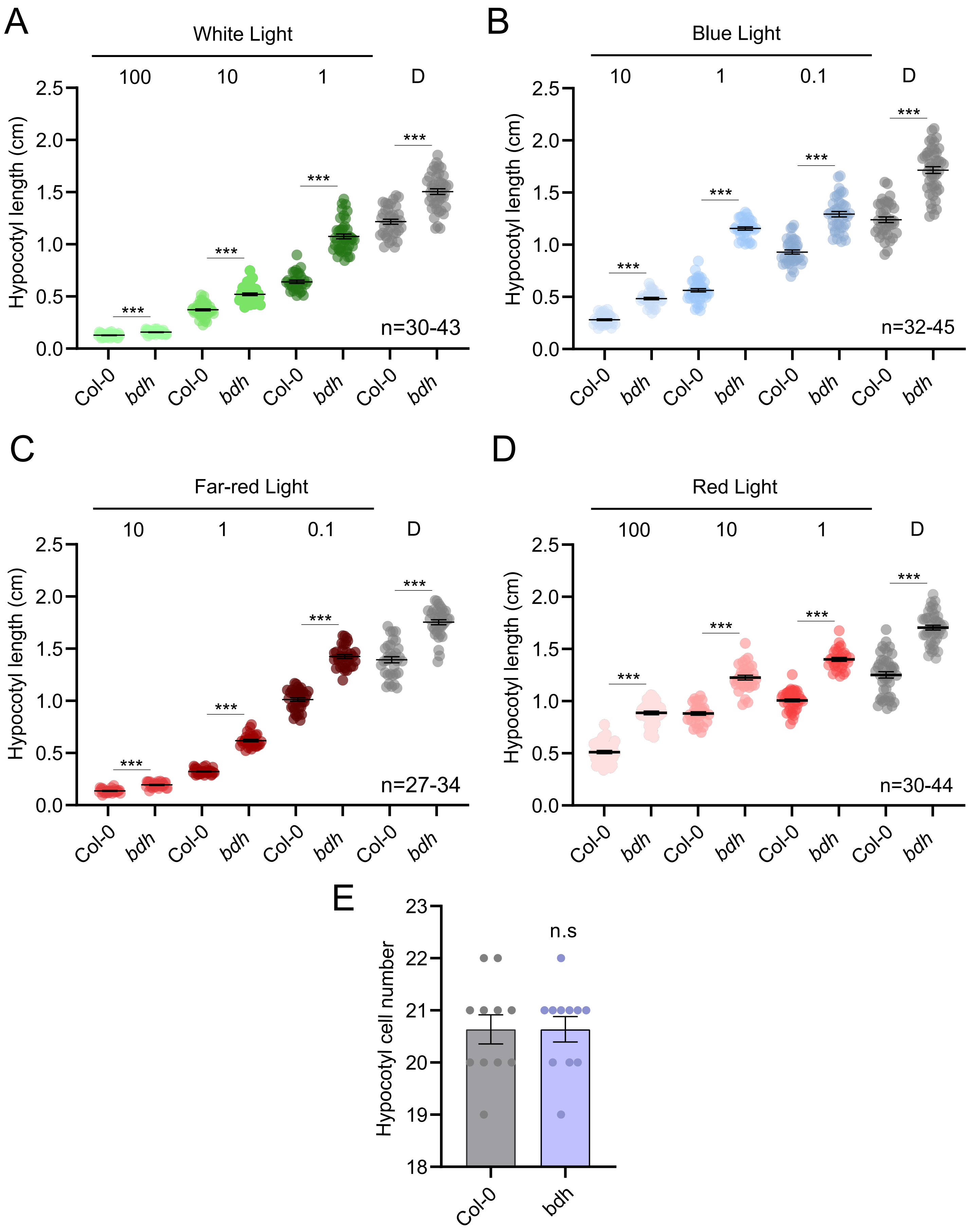


**Supplementary Figure 2. BDH regulate hypocotyl growth under different light qualities**

(A-D) Dot plots depicting hypocotyl length of 7-day-old Col-0 and *bdh* mutants under a series of white (A), blue (B), far-red (C), and red (D) light intensities measured as μmol $m^{-2}$ $s^{-1}$ as indicated in the upper region of each panel. D: darkness. Each dot represents hypocotyl length of one individual seedlings . Range of seedlings numbers (n) used in each experiment are shown in each panel. Asterisks indicate significant differences (P < 0.05), as determined by Student's t-test. Error bars represent Mean ± SEM, with n=27-45. At least two independent biological replicates were conducted with similar results. (E) Plot depicting hypocotyl cell number of 7-day-old seedlings grown in long-days condition. Each dot represents the total number of cells measure in one individual hypocotyl of WT and *bdh* mutant. Error bars represent Mean ± SEM, with n=10. Three independent biological replicates were conducted with similar results. Student's t-test, *P < 0.05. n.s, not significant.

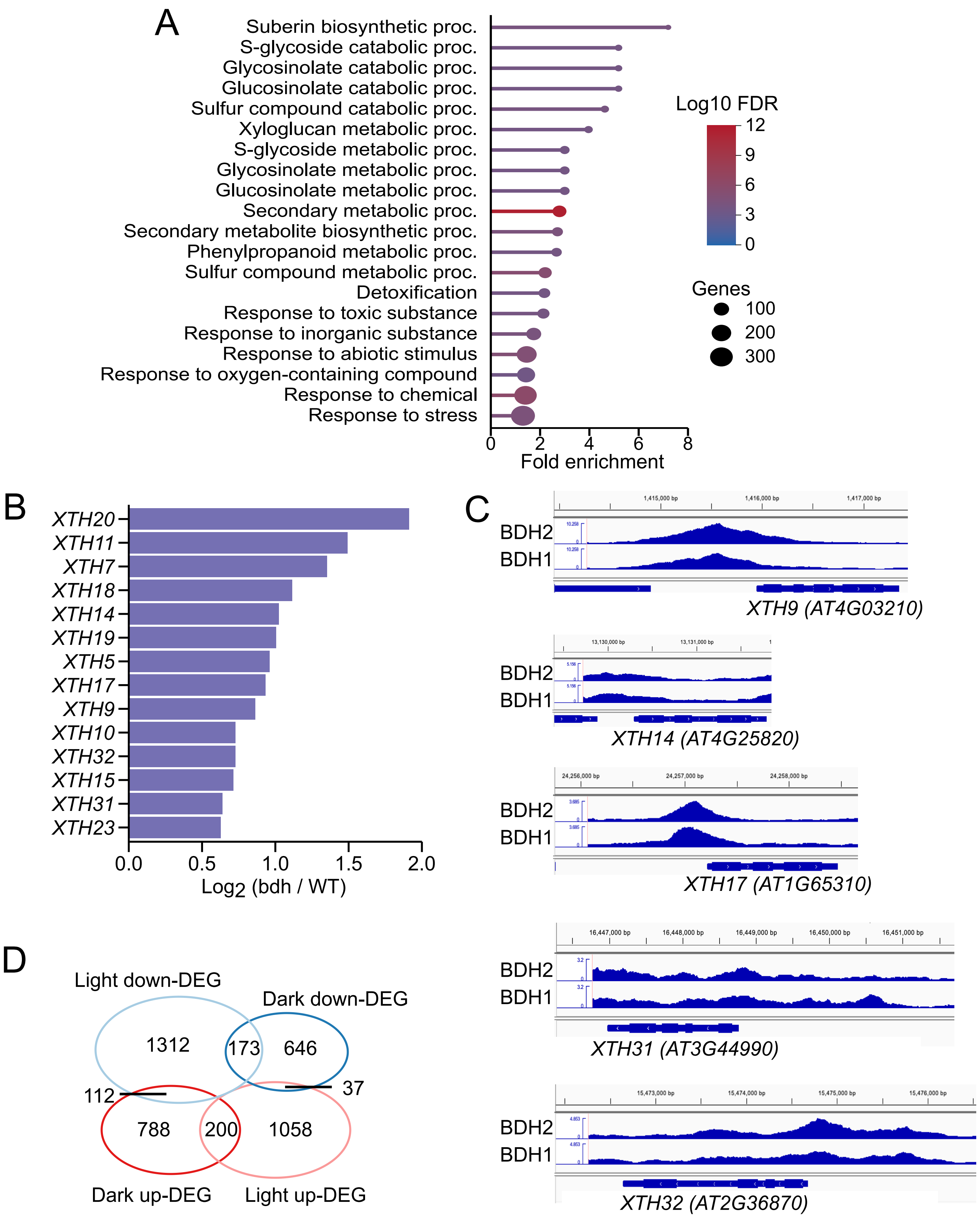


**Supplementary Figure 3. BDH regulates the expression of *XTH* genes**

(A) Dot-plot showing GO enrichment analysis performed over bdh DEGs detected in 5-day-old seedlings grown in darkness. The top 20 most significant categories are represented. (B) Log2 fold changes (FC) (bdh/WT) of XTH genes that were classified as DEGs in 5-day-old *bdh* etiolated seedlings (log2FC 0.58, q-value 0.05) (C) Browser screenshots of a published BDH1 and BDH2 ChIP-seq in light-grown seedlings[10] depicting ChIP-seq signal over *XTH9, XTH14, XTH17, XTH31* and *XTH32* genes, selected as representative BDH-regulated genes in both dark and light conditions (Figure 1D). (D) Overlap between *bdh* upregulated and downregulated DEGs detected in darkness and light. The RNA-seq data from light-grown seedlings was obtained from[6].

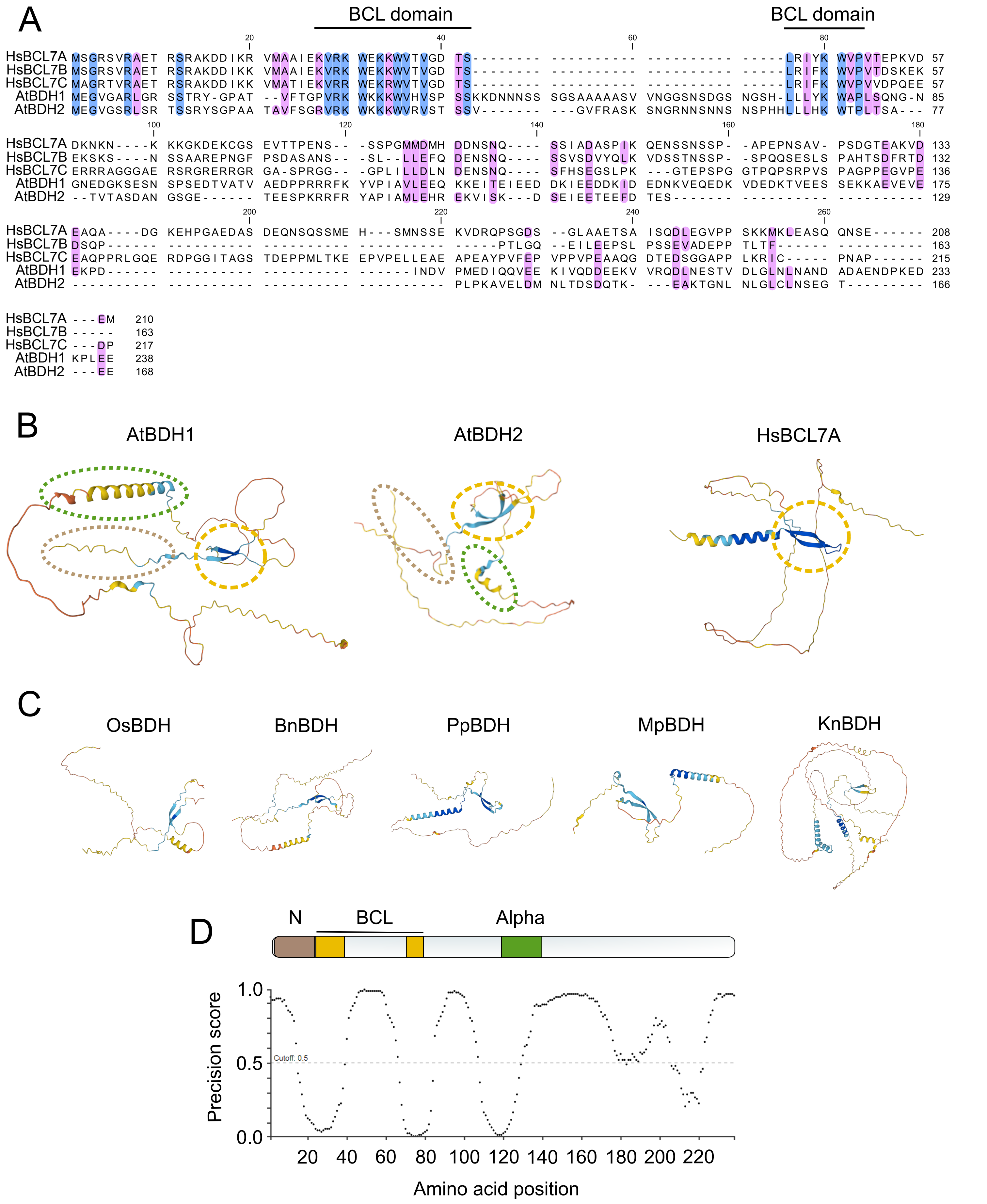


**Supplementary Figure 4. Analysis of conservation of BDH domains.**

(A) Sequence alignment of *Homo sapiens* (*Hs*) BCL7A, BCL7B, and BCL7C, and *Arabidopsis thaliana* (*At*) BDH1 and BDH2. Conserved residues in at least four proteins are highlighted in blue, while similar residues in at least four proteins are shaded in pink. The BCL domain is indicated at the top of the alignment according to[13] (B) AlphaFold 2 models depicting the structure prediction of AtBDH1, AtBDH2, and HsBCL7A. The N-terminal domain region is highlighted in brown, the BCL domain's β-hairpin in orange, and the alpha helix of the Alpha domain in green. (C) AlphaFold 2 models depicting the structure prediction of BDH orthologs from various plant species: *Oryza Sativa* (*OsBDH*; A0A0P0XL11), *Brassica napus* (*BnBDH*; A0A816N4Q3), *Physcomitrella patens* (*PpBDH*; A0A2K1ICB9), *Marchantia polymorpha* (*MpBDH*; A0A2R6X4Y7), *Klebsormidium nitens* (*KnBDH*; A0A1Y1IQI7). (D) Schematic representation of the Arabidopsis BDH1 protein, highlighting its N-terminal (brown), BCL (orange), and Alpha domains (green). The accompanying graph depicts the prediction of disordered regions within the protein obtained from to DISOPRED3[30].

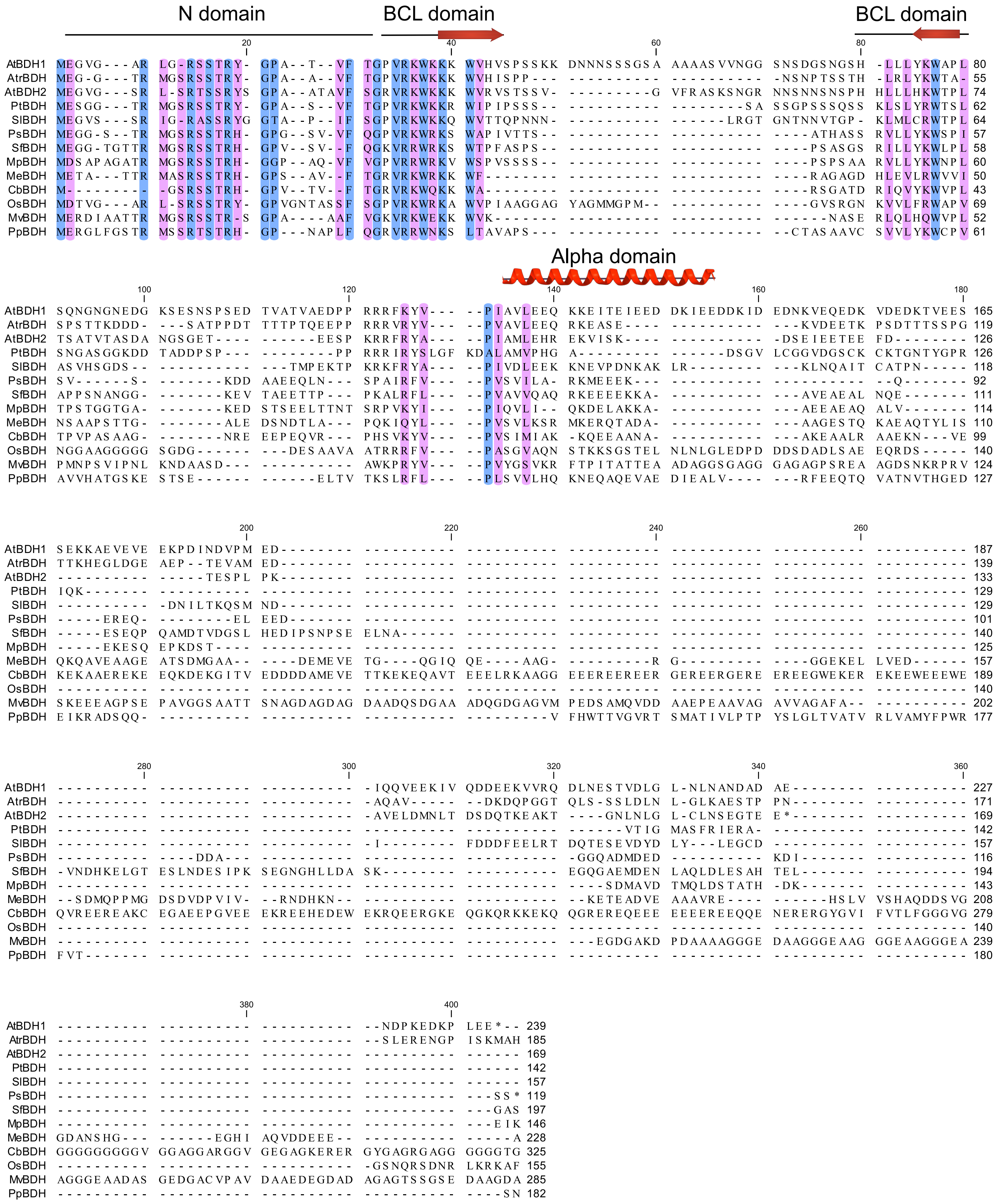


**Supplementary Figure 5. Evolutionary conservation of BDH across plants.**

Alignment of BDH protein sequences across Viridiplantae, including the following species: *Arabidopsis thaliana, Amborella trichopoda, Chara braunii, Marchantia polymorpha, Mesostigma viride, Mesotaenium endlicherianum, Oryza sativa, Physcomitrium patens, Pinus taeda, Selaginella moellendorffii, Solanum lycopersicum, and Sphaerobunus fallax*. Conserved residues in at least eleven proteins are highlighted in blue, while similar residues in at least eleven proteins are shaded in pink. The regions corresponding to the characterized domains (N, BCL, Alpha) are indicated at the top of the alignment, based on both conservation and structure prediction using AlphaFold 2 (Supplementary Figure 4).

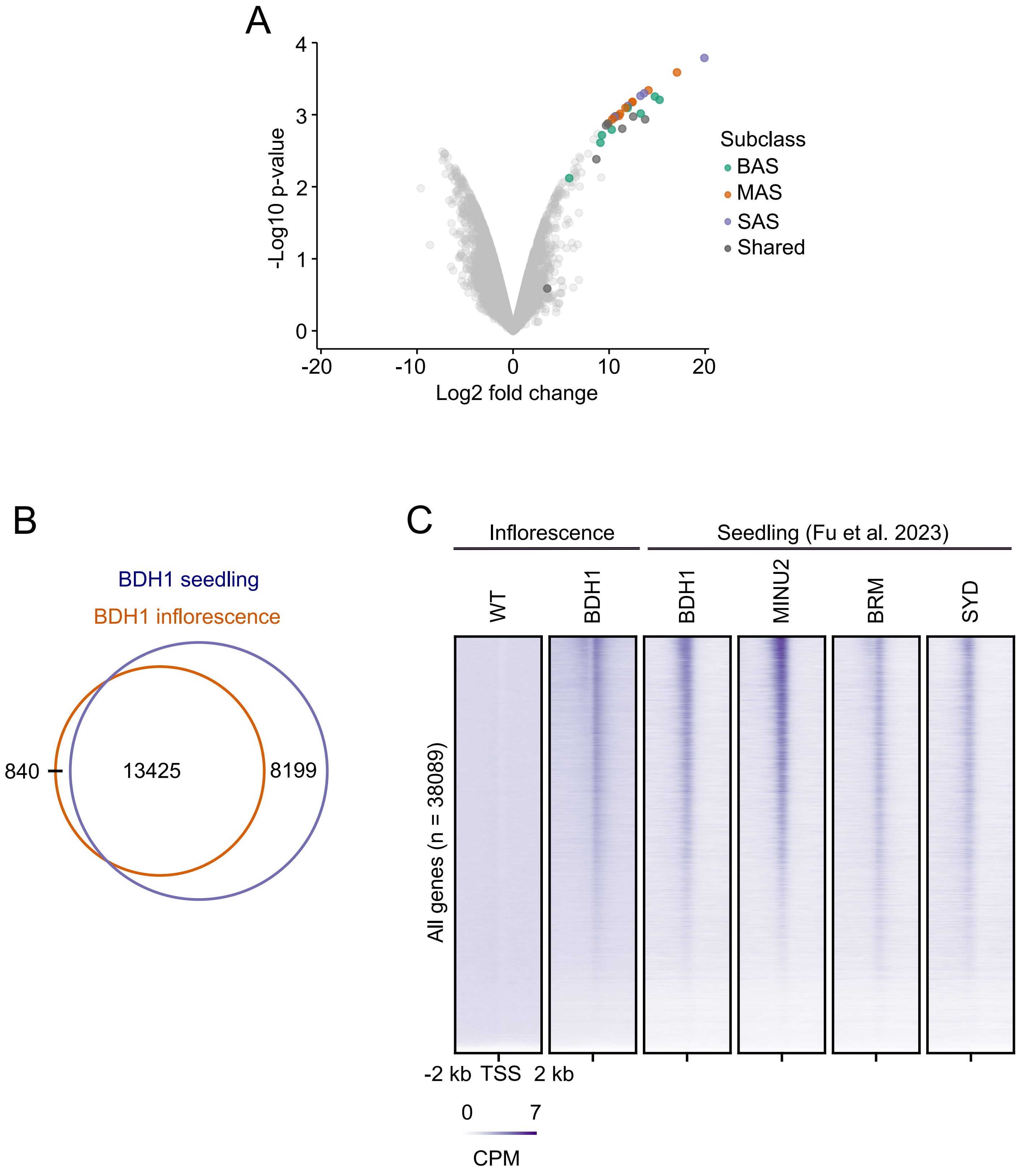


**Supplementary Figure 6. Protein interactome and genomic targets of BDH1 in inflorescences.**

(A) Volcano plot depicting enrichment of proteins immunoprecipitated with BDH1-3xFLAG in *bdh1* single mutant background. Distinct colours represent SWI/SNF subclass-specific subunits. X axis depicts log2 fold change of average intensities of IP experiments. Y axis depicts significance -log10 p-value. (B) Overlap between BDH1 ChIP-seq peaks in inflorescences and seedlings. Genes that either overlapped peaks or whose TSS or TES was less than 2kb away from a peak summit were considered as targets. (C) Heatmap showing the accumulation of non-transgenic Col-0 control (WT) and BDH1 in inflorescences and BDH1, MINU2, BRM, and SYD in seedlings over the TSS of Araport11 genes (n = 38089). SWI/SNF ChIP-seq data in seedlings was obtained from[10].

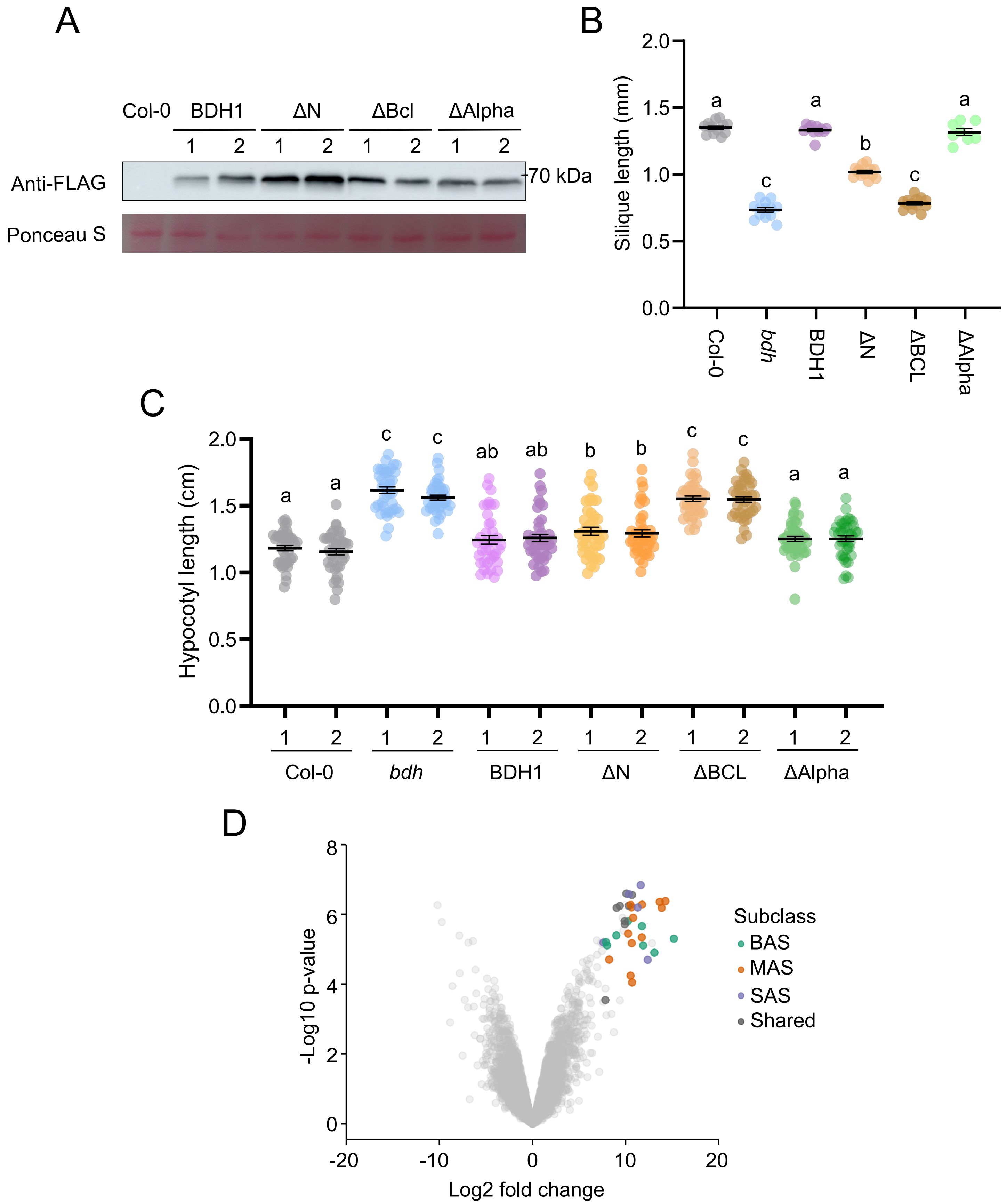


**Supplementary Figure 7. Functional characterization of BDH domains**

(A) WB analysis depicting the protein accumulation in two independent transgenic lines expressing 3xFLAG-tagged full length BDH1, or the following depletions: ΔN, ΔBCL, and ΔAlpha. Ponceau S staining of Rubisco is shown as loading control.
(B) Dotplot depicting the silique length (mm) in the primary inflorescence of Col-0, *bdh* mutant, and independent T1 plants expressing BDH1, ΔN, ΔBCL, and ΔAlpha. Error bars represent Mean ± SEM, with n=8-13. (C) Dotplot depicting the hypocotyl length (cm) of T2 populations of 7-day-old etiolated seedlings of the labelled backgrounds. Error bars represent Mean ± SEM n=40-44. (B-C) Different letters indicate significant differences ($P < 0.05$), as determined by ANOVA with Tukey's post-hoc test.
(D) Volcano plot depicting enrichment of proteins immunoprecipitated with BDH1-3xFLAG in *bdh* mutant background. Distinct colours represent SWI/SNF subclass-specific subunits. X axis depicts log2 fold change of average intensities of IP experiments. Y axis depicts significance -log10 p-value.

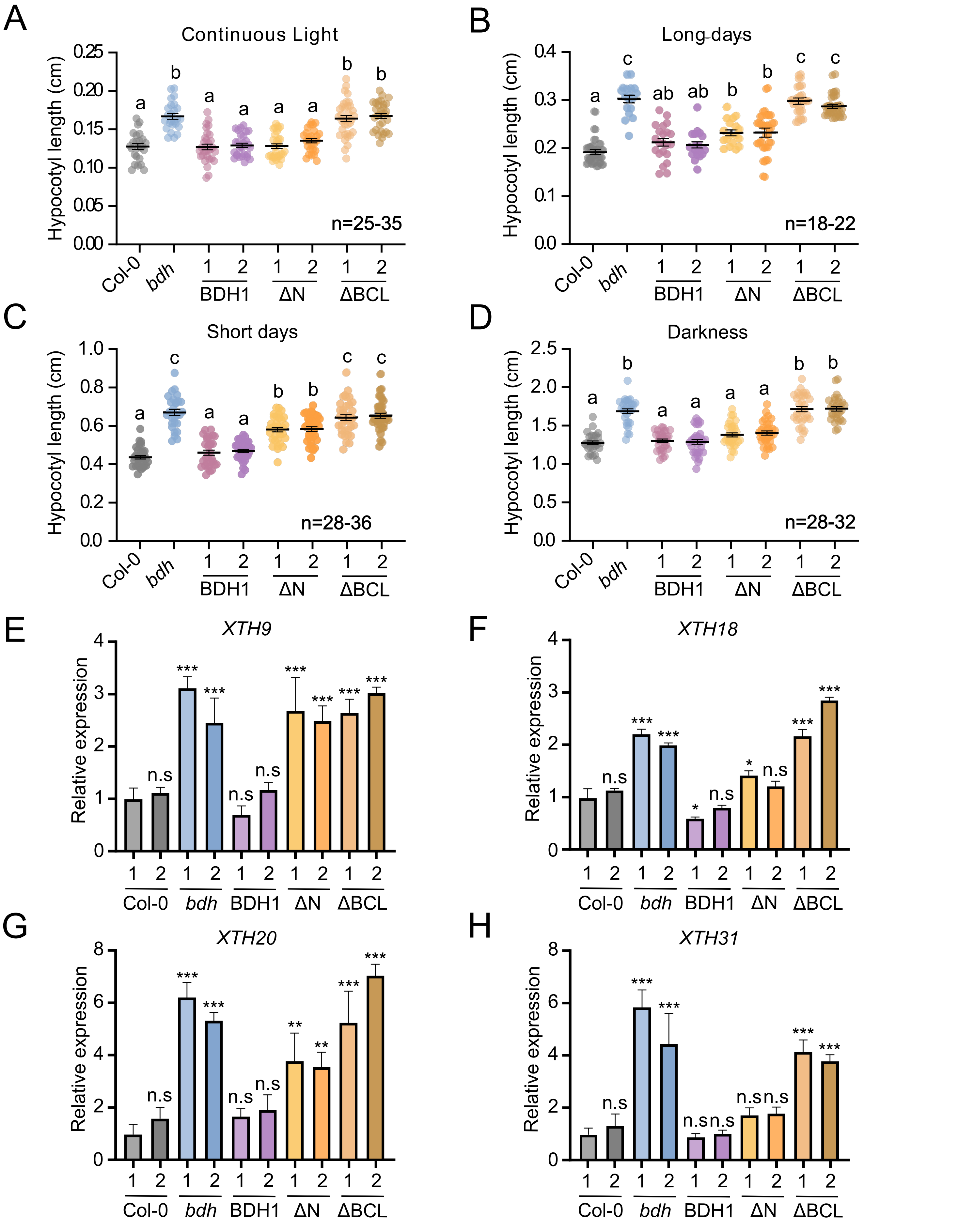


**Supplementary Figure 8. Characterization of BDH domains function in etiolated plants**

(A-D) Dot plots depicting hypocotyl length (cm) of 7-day-old Col-0, *bdh*, and two independent T3 generation lines of BDH1, ΔN, and ΔBCL under (A) continuous light, (B) long days, (C) short days, and (D) darkness (n=20-35). Each dot represents hypocotyl length of one individual seedlings . Range of seedlings numbers (n) used in each experiment are shown in each panel. Error bars represent Mean ± SEM. Different letters indicate significant differences (P < 0.05), as determined by ANOVA with Tukey's post-hoc test. Two independent biological replicates were conducted with similar results. (E-H) RT-qPCR results showing the expression of (E) *XTH9*, (F) *XTH18*, (G) *XTH20*, and (H) *XTH31* in 5-day-old etiolated seedlings of Col-0, bdh, and two independent T3 generation lines of BDH1, ΔN, and ΔBCL. Results were normalized against the housekeeping gene PP2A using the ΔΔCt method. Asterisks denote statistically significant differences between mean values, as assessed by Student's t-test (*P < 0.05, **P < 0.01, ***P < 0.001, n.s., not significant).

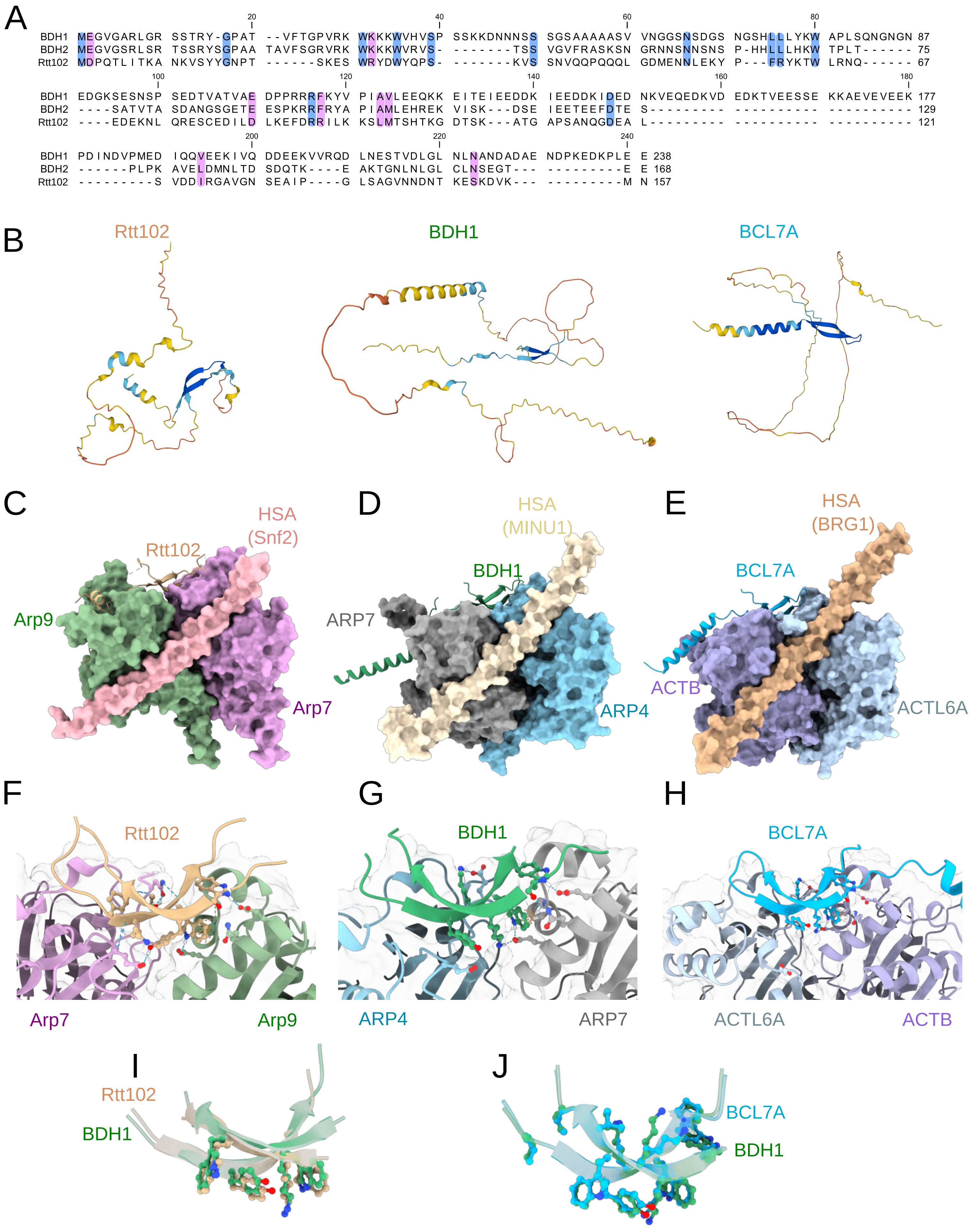


**Supplementary Figure 9. Conservation of the SWI/SNF catalytic module across eukaryotes.**

(A) Sequence alignment of *Saccharomyces cerevisiae* Rtt102 and *Arabidopsis thaliana* BDH1 and BDH2. Conserved residues in the three proteins are highlighted in blue, while similar residues are shaded in pink. (B) AlphaFold2 structural models of BDH1, BDH2, and Rtt102 are depicted. (C-E) Yeast with experimental structure (PDB:4I6M) along structural models of Plant (D) and Human (E) complexes. The conserved interaction of the β-hairpin and ARPs is shown in (F-H). Hydrogen bonds and hydrophobic interactions are highlighted in blue and orange dashed lines respectively. (I-J) Detail of the structural alignment of Arabidopsis BDH1 and (I) Yeast Rtt102 or (J) Human BCL7A structures. The side chains of conserved amino acids are shown. Arabidopsis model (green), Yeast structure (PDB 4I6M) (brown) and Human model (blue). Aminoacids are colored by heteroatom (red: oxygen; dark blue: nitrogen).

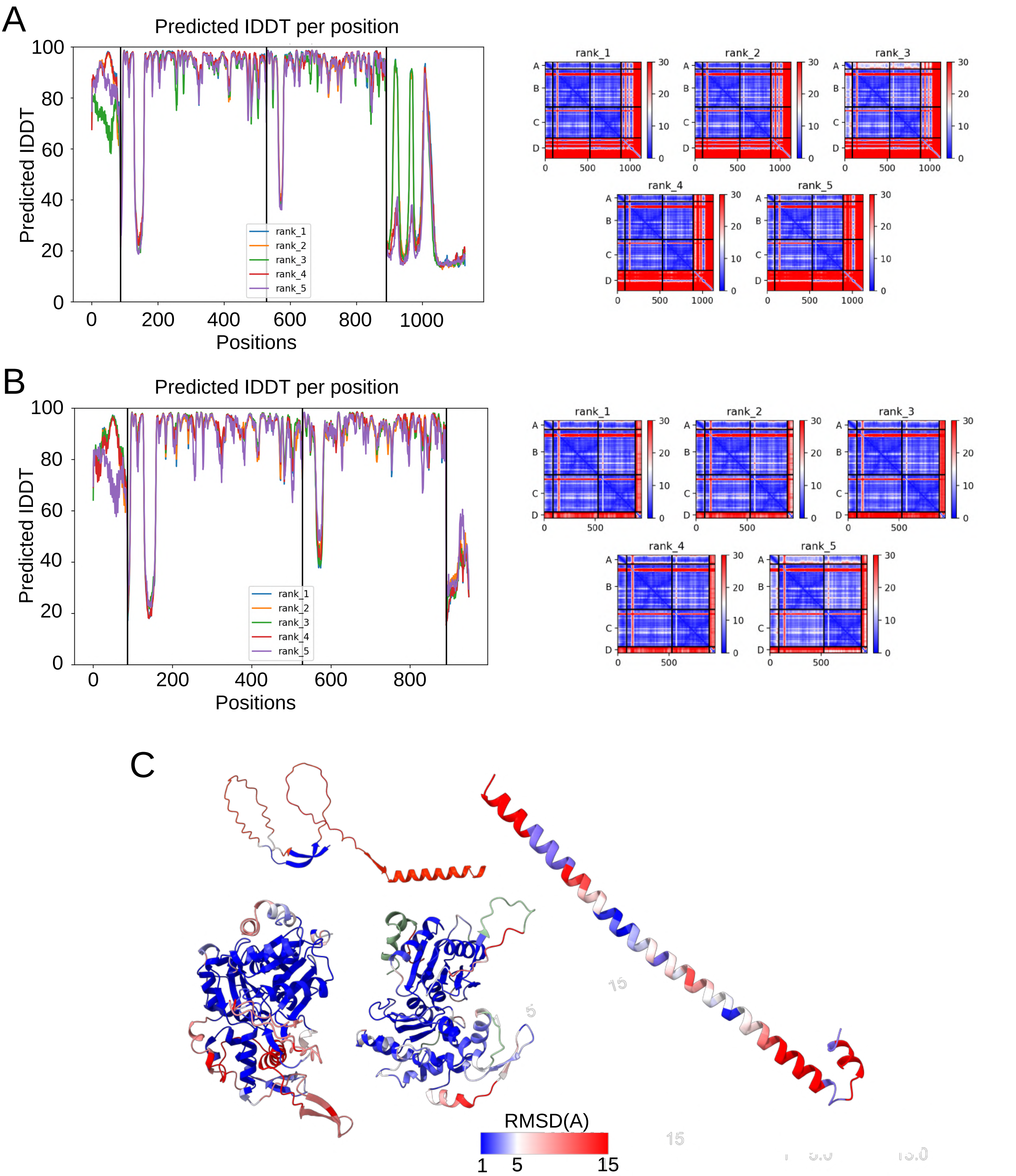


**Supplementary Figure 10. Analysis of the Alphafold2-multimer complex predictions**

(A) Predicted total distance difference test (pLDDT) values (left) and predicted aligned error in Å (right) for Arabidopsis MINU1$^{(HSA)}$-ARP4-ARP7-BDH1 predicted complex. (B) Predicted total distance difference test (pLDDT) values (left) and predicted aligned error in Å (right) for Human BRG1$^{(HSA)}$-ACTBL6-ACTBL-BCL7 predicted complex. (C) Structural alignment of PDB 4I6M (yeast, Arp7-Arp9-Snf2$^{(HSA)}$-Rtt102) to the Arabidopsis model (ARP4-ARP7-BDH1-MINU1$^{(HSA)}$). The root mean square deviation (RMSD) of the predicted structure compared to the experimental structure PDB 4I6M is shown. RMSD is represented in Amgstrongs.

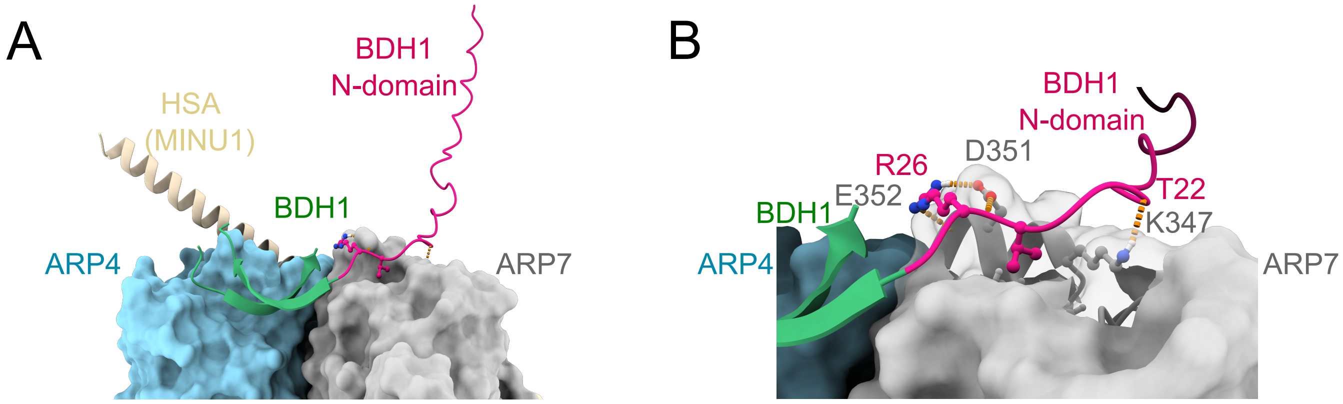


**Supplementary Figure 11. Structural prediction of BDH1 N-term in the catalytic module**

(A) Overview and (B) detail of the BDH1 N-terminal (pink) (BDH1 aminoacids 1-27). Hydrogen bonds are depicted in orange dashed lines.

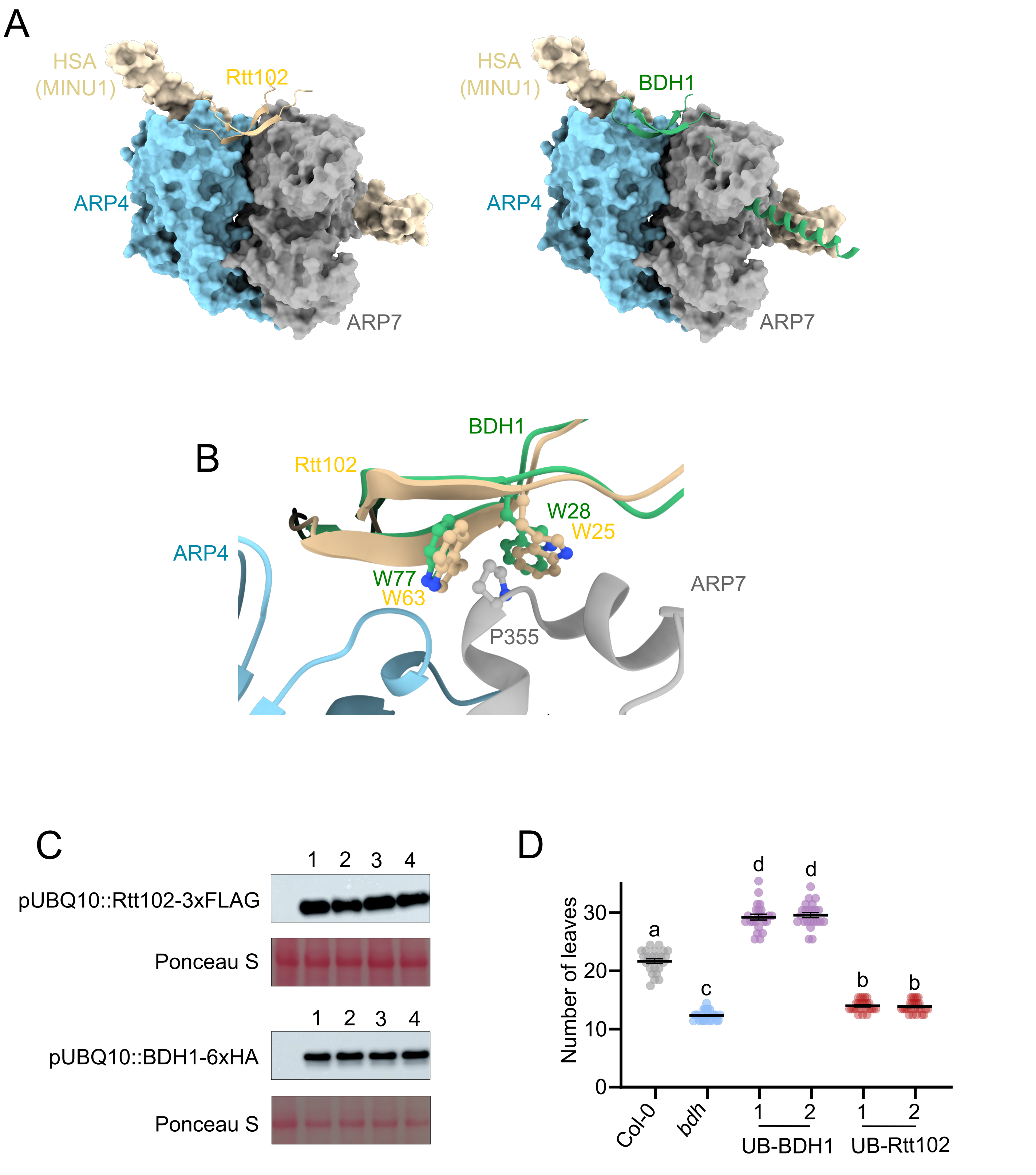


**Supplementary Figure 12. Structural and functional conservation of Rtt102 and BDH proteins**

(A) Structural model predicting the interaction of the yeast protein Rtt102 with the catalytic module of the plant complex (left), compared to the plant complex including BDH1(right). (B) Structural alignment of BDH1 and Rtt102 β-hairpins in complex with ARP4-ARP7 heterodimer. The predicted interaction between conserved tryptophans in BDH1 and Rtt102 and Arabidopsis ARP7 P355 are shown. (C) WB analyses depicting the protein accumulation levels of four independent T1 transgenic lines of pUBQ10::Rtt102-3xFLAG (UB-Rtt102; top panel) and pUBQ10::BDH1-6xHA (UB-BDH1; bottom panel) in the *bdh* background. Ponceau S staining of Rubisco is shown as loading control. (D) Dotplot depicting flowering time measured as the total number of rosette and caulinar leaves after bolting in plant grown in long-days conditions of Col-0 and *bdh* backgrounds, and two independent T2 populations of plants expressing pUBQ10::BDH1-6xHA (UB-BDH1) and pUBQ10::Rtt102-3xFLAG (UB-Rtt102). Different letters indicate significant differences ($P < 0.05$), as determined by ANOVA with Tukey's post-hoc test. Error bars represent Mean ± SEM n=21-25.

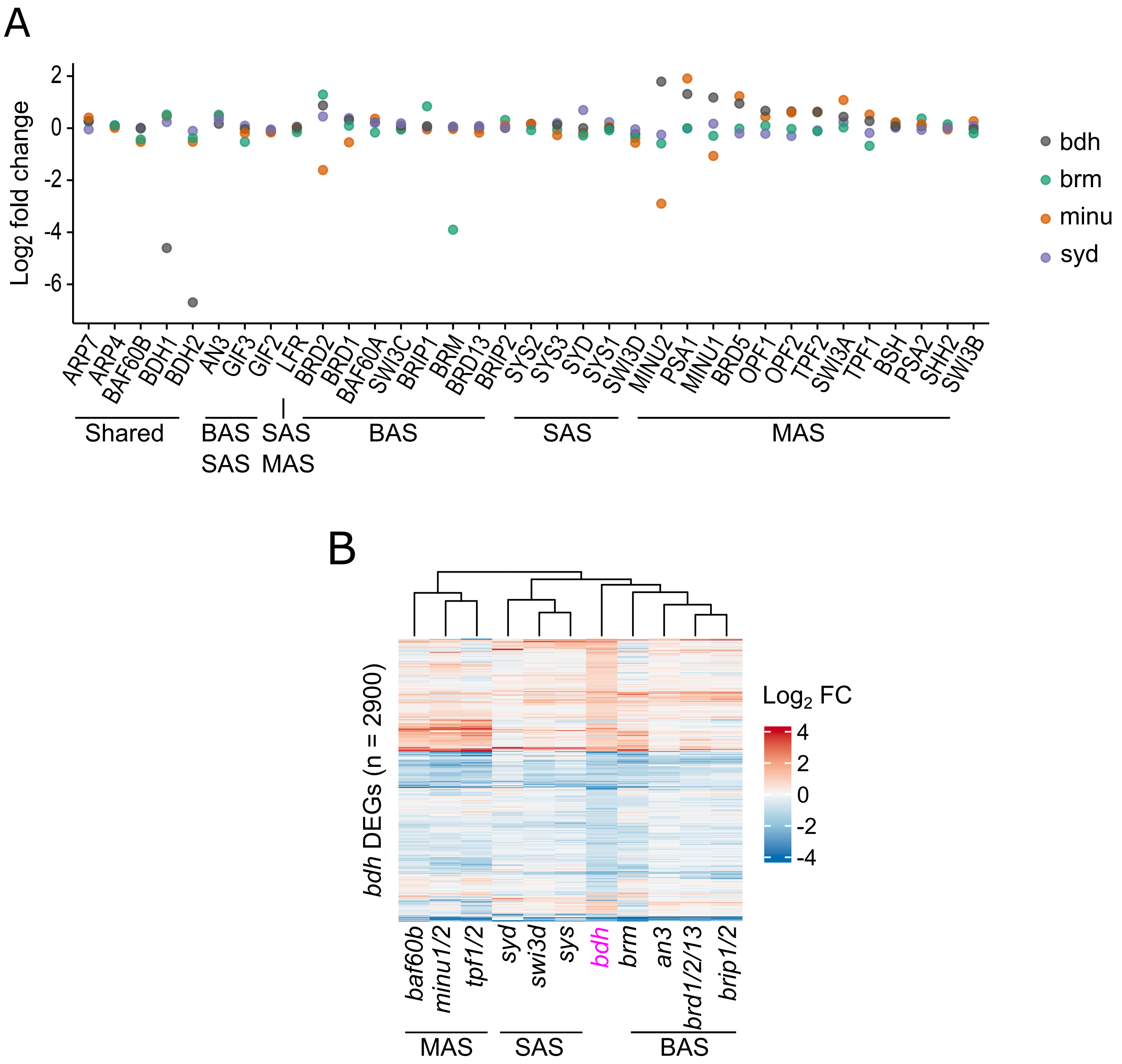


**Supplementary Figure 13. BDH stabilises ARP heterodimer and functions in all SWI/SNF subclasses**

(A) Dot plot depicting expression changes of SWI/SNF subunits in the *bdh* (grey), *brm* (green), *minu* (orange), and *syd* (blue) mutant backgrounds. The MAS, SAS and BAS subclass-specific subunits, and those shared by more than one subclass are indicated. RNA-seq data from[9]. (C) Heatmap showing hierarchical clustering of the expression changes of SWI/SNF mutants over bdh DEGs. RNA-seq data from[9]. The MAS, SAS and BAS subclass-specific mutants are indicated. The bdh mutant is highlighted in pink .

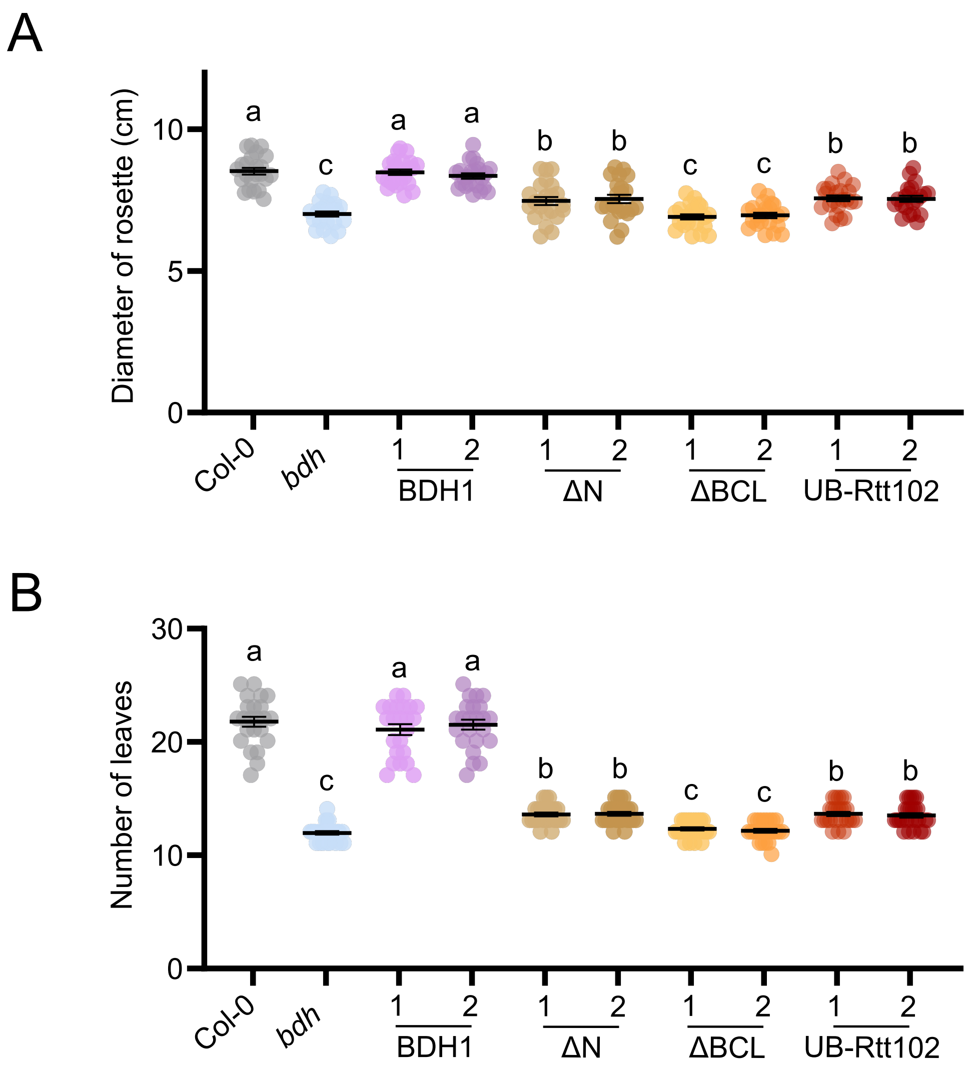


**Supplementary Figure 14. Rtt102 rescues the *bdh* mutant to the same extent as N-term deleted BDH1**

(A) Length of the widest measured rosette diameter in 28-day-old Col-0, *bdh*, and two independent T2 populations from BDH1, ΔN, ΔBCL and UB-Rtt102 lines under long-day conditions. (B) Number of total leaves after bolting in the labelled backgrounds under long-day conditions. The different letters indicate significant differences ($P < 0.05$), as determined by ANOVA with Tukey's post-hoc test. Error bars represent Mean ± SEM, n =22-23.
